# Ensemble of BLUP, Machine Learning, and Deep Learning Models Predict Maize Yield Better Than Each Model Alone

**DOI:** 10.1101/2023.03.30.532932

**Authors:** Daniel R. Kick, Jacob D. Washburn

## Abstract

Predicting phenotypes accurately from genomic, environment, and management factors is key to accelerating the development of novel cultivars with desirable traits. Inclusion of management and environmental factors enables *in silico* studies to predict the effect of specific management interventions or future climates. Despite the value such models would confer, much work remains to improve the accuracy of phenotypic predictions. Rather than advocate for a single specific modeling strategy, here we demonstrate within large multi-environment and multi-genotype maize trials that combining predictions from disparate models using simple ensemble approaches most often results in better accuracy than using any one of the models on their own. We investigated various ensemble combinations of different model types, model numbers, and model weighting schemes to determine the accuracy of each.

We find that ensembling generally improves performance even when combining only two models. The number and type of models included alter accuracy with improvements diminishing as the number of models included increases. Using a genetic algorithm to optimize ensemble composition reveals that, when weighted by the inverse of each model’s expected error, using combinations of best linear unbiased predictors, linear fixed effects models, deep learning models, and select machine learning models perform best on our datasets.

## Introduction

Phenotypic prediction based on environmental, management, and genetic factors would support improvements in crop yield through model driven selection and yield forecasting. In both cases, the accuracy of a given model is key for the model to inform decisions: inaccurate and misleading models could result in substantial repercussions in terms of time and resources.

Over the past few decades, increasing efforts have focused on the development of predictive models for genomic selection. While incorporating substantial amounts of environmental information into these models has not been common historically, this has recently become an area of increasing interest. The combination of genomic and environmental effects (both those within a farmer’s control, i.e., management, and those external to it) are necessary because gene by environmental effects can in some cases exceed purely genetic effects (Rogers et al. 2021). Furthermore, inclusion of these interactions can improve the predictive accuracy for new environments or cultivars (Li et al. 2021; Jarquin et al. 2021; Montesinos-López et al. 2023).

Efforts to improve phenotypic prediction in agriculture have also involved the development of mechanistic physiological crop growth models (Technow et al. 2015) and data driven models – both statistical (Jarquin et al. 2021; Rogers et al. 2021; Rogers and Holland 2021) and machine learning models (Westhues et al. 2021; Washburn et al. 2021). Combining physiological and statistical models into a single model has also been explored but not yet widely implemented (Messina et al. 2018; Diepenbrock et al. 2021; Shahhosseini et al. 2021a). Among models with substantial environmental, as well as genetic, components, Best Linear Unbiased Predictor models (BLUPs) and deep neural networks (DNNs) have both been shown to perform well under some scenarios (Washburn et al. 2021) with a potential tradeoff in average model performance favoring BLUPs and model consistency across replicates in some cases favoring DNNs (Kick et al. 2023) in addition to the ability to more directly incorporate multimodal data (Montesinos-López et al. 2023). For most breeding applications however, BLUP models with limited or no environmental data are still considered the gold standard with their relative simplicity, when compared to machine learning, deep learning, and other approaches being a major benefit (Ma et al. 2018; Montesinos-López et al. 2018; Abdollahi-Arpanahi et al. 2020; Nazzicari and Biscarini 2022; Gianola et al. 2022).

While extensive attention has been given to which modeling strategies are expected to produce the best results for phenotypic prediction, much less consideration has been given to using predictions from multiple models simultaneously. This is particularly true in the context of models that include genetic, environment, and/or management data as opposed to the standard genomic selection paradigm using only genomic data (Banerjee et al. 2020; Abdollahi-Arpanahi et al. 2020). Ensemble learning is a method by which the predictions between multiple models – ideally from a diverse set of models are aggregated (Zhou 2015). This results in predictions which are frequently superior to those from the base models. Methods for creating ensembles include averaging predictions using uniform or varied weights and training a model using the predicted values as inputs – “stacking” a model on the base models – among other approaches. Certain common machine learning methods such as random forests, operate by ensembling many weakly informative models (in this case decision trees). Here we use ensemble learning to refer to aggregations of models which could have been used individually (e.g., combining a BLUP and random forest) rather than those which *require* ensembling to function (e.g., a single random forest model).

Ensemble learning has been shown to be effective in as varied uses as forecasting solar and wind availability (Carneiro et al. 2022) and optimizing genetic transfer in chrysanthemum using *Agrobacterium* (Hesami et al. 2020) to, in some cases, genomic prediction of phenotypes (Azodi et al. 2019; Liang et al. 2021). In addition to the examples above, ensemble methods have been used within agriculture to predict soybean yield based on measured yield component traits (Yoosefzadeh-Najafabadi et al. 2021b). They have also been applied to making predictions using optical (e.g. hyperspectral) data in alfalfa (Feng et al. 2020), winter wheat (Li et al. 2022) and soybean (Yoosefzadeh-Najafabadi et al. 2021a). Within maize, ensembling has been found to be effective for models predicting yield at a high level – predicting yield of counties rather than plots – using machine learning (Shahhosseini et al. 2020), machine learning coupled with crop modeling (Sajid et al. 2022), and deep learning (Shahhosseini et al. 2021b).

Ensemble learning has been found to be beneficial for prediction in a study considering soy, maize, rice, sorghum, switchgrass, and spruce datasets (Azodi et al. 2019). Similar results have been seen in genomic selection models of Chinese Simmental beef cattle (Liang et al. 2021).

Notably, these studies focused on the influence of genetics without consideration of GxE effects. Incorporating predictions from an ensemble model to account for nonlinear effects has been found useful in capturing GxE effects in wheat (Heslot et al. 2014). In this case the ensemble model was incorporated into a linear model rather than being the final predictive model. Despite these examples demonstrating that ensemble learning can be effective, use in conjunction with modeling environmental interactions has not gained wide adoption for phenotypic prediction in crops, and is relatively understudied when compared to non-ensemble approaches.

If effective for complex models containing genomic and environmental components, phenotype prediction and selection could be immediately improved merely through utilizing combinations of commonly applied (or less common) models that are often already being created by researchers or breeders for the purpose of benchmarking. The aim of this study is to establish whether this is the case in large maize data sets with extensive genomic, environmental, and management data. An additional goal is to identify the types and numbers of models, along with the specific ensembling strategies, that are most effective at improving phenotypic prediction. To accomplish this, diverse maize yield prediction models are used to demonstrate that ensembling is an effective means to decrease error in a simple case where only two individual models are combined. In fact, we show that within the tested datasets and methods there is not a case when one of the ensemble models results in higher expected error than both of its parent models. Next, we consider the utility of increasing the number and variety of models used and find that increasing the number of models generally increases performance, but with diminishing returns as more and more models are added. Finally, we consider all combinations of multiple models, more than two model types, and different weighting schemes for aggregating predictions. Based on optimization with a genetic algorithm, aggregation of predictions from best linear unbiased predictors, consecutively optimized deep neural networks, linear fixed effects models, and select machine learning models (support vector regression and random forest) with predictions weighted inversely proportionate to expected model error was optimal for these data.

## Materials and Methods

### Data Preparation

Maize yield, environmental, and management data came from the Genomes to Fields (G2F) initiative’s data releases for 2014-2019 (McFarland et al. 2020). These data are publicly available (https://www.genomes2fields.org/resources/) and provide weather and soil data for fields in the continental United States, management information, and genomic data in addition to phenotypic measurements. Weather data was supplemented with data from Daymet (Thornton et al. 2020). Genomic, environmental, and management data was quality controlled using custom scripts and released previously through Zenodo (cleaned data 10.5281/zenodo.6916775 and scripts 10.5281/zenodo.7401113) accompanying a previous study (Kick et al. 2023).

### Statistical and Machine Learning Models

Using phenotypic models, yield was predicted from genomic, environmental, and management data. While all models considered here used data from these categories, they vary in the representation of data used and model complexity. As processing of predictions subsequent to their generation is the focus of this analysis, a brief summary is provided here with full details being available (Kick et al. 2023). The considered models included linear models, machine learning models, and deep learning models. Among linear models, linear fixed effects model (LM), and a best linear unbiased predictor model (BLUP) were considered. Within machine learning models four model types were considered: k nearest neighbors (KNN), radius neighbor regression (RNR), support vector regression (SVR), and random forest regression (RF). Lastly, two deep neural networks are considered. The first neural network resulted from “consecutive optimization” of subnetworks predicting yield from genomic, soil, weather and management, and the interactions between these data types (DNN-CO). The second resulted from “simultaneous optimization” of all subnetworks at once (DNN-SO).

### Assessing Model Performance

To assess model performance available data was split into a training and testing set with the former being used to fit models and the latter being used to evaluate model accuracy. These sets were constrained such that no observations from the same field and year were allowed in both sets. This requires models to predict yield in a new growing season which increases the difficulty and value of generating accurate predictions. For a set of predicted yields, accuracy was quantified using root mean squared error 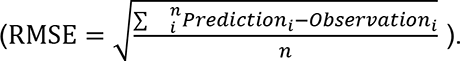 models are sensitive to randomly initialized values and thus replicates of the same model are not guaranteed to produce identical predictions. To account for this variability, 10 replicate models were trained and used.

### Ensembling Methods

Model predictions were combined using five methods. The first method was an arithmetic mean (uniformly weighting models) of the predictions. The second, third, and fourth, methods were weighted averages with weights inversely proportionate to the number of like models included (i.e., weighted such the total weight for each model type is equal), the standard deviation of each set of predictions, and the variance of each set of predictions. The fourth method weighed predictions from models inversely proportionate to each model’s RMSE based on the training data (using RMSE on the test data would be an inaccurate assessment of this method). The final method trained a small multilayer perceptron with two layers consisting of 100 neurons each to predict yield based on 80 predicted yield values (10 replicates of the 8 models listed above). This model was trained for 500 epochs using scikit-learn’s (Pedregosa et al. 2011) defaults.

### Evaluating Composition of Model Ensembles

While calculating the RMSE of all possible pairwise model averages is trivial, calculating the RMSE of all combinations of our model replicates, even with a single ensembling strategy is not (1.2x10^24^ combinations). Instead, a Monte Carlo simulation approach was used to approximate the distribution RMSEs resulting from a given ensembling strategy rather than exhaustive calculation. To determine effective strategies for using multiple models several simplified cases were considered followed by an optimization approach. First, ensembling two model replicates of the same or differing model types was considered using an arithmetic mean to combine the predictions. Second, the effect of increasing the number of models used was considered along with different selection schemes – selected at random, equally across all model types, or using the pairings initially considered – and several weighting approaches.

Finally, analysis was expanded to include different numbers of models from each model type by using a genetic algorithm to identify the optimal combination and ensembling method.

### Ensembles Composed of Two Models

Initially the performance of all combinations of model replicates when their predictions were averaged was evaluated. These were compared against each other and the performance of the base models. The likelihood of an ensemble improving performance relative to the average base model was calculated. Analysis of other weighting schemes was considered as part of following analyses.

### Ensembles Composed of Variables Models of One or More Model Types

Building off the pairwise comparisons above, the expected RMSE was simulated from a given combination and number of models. This procedure was consistent for the combinations explored below. For a given number of models for each model type and model ensembling method (except the multilayer perceptron-based method) a set of 50 RMSEs were simulated.

The target number of model replicates for each model type were selected randomly with replacement then ensembled and scored. The average of the simulated distribution was taken as the expected RMSE.

After simulating the expected RMSEs over a range of models included (i.e., either the total number of models used or the models of each model type used) a function was fit to describe the trend. As some model replicates do not have variation between their predictions (i.e., model replicates converged or effectively converged), increasing the number of models used will not necessarily alter ensemble performance. These combinations were treated as a linear model and identified by the standard deviation of all collected RMSEs being less than or equal to 1x10^-3^. For all other cases a three-term exponential decay model 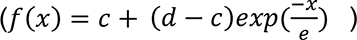) was fit using the library “drc” (Ritz et al. 2015). Note that here “e” is a variable.

The model terms “c” and “d” were allowed to vary for each combination of model types. As a result, the effect of increasing the number of models being ensembled was allowed to saturate which was observed in certain exploratory visualizations.

This procedure was applied to assess the effect of increasing the number of ensembled models in three cases. First, ensembling models selected at random with respect to model type were considered. Between 1 and 80 models were selected to be used. This deviates from the procedure outlined above by simulating 50 combinations once at each step rather than one combination 50 times. Next, a uniform mix of all model types was assessed, selecting between 1 and 10 replicates of each model for the simulated ensemble. Finally, pairwise combinations of model types (including a model type combined with its own replicates) were simulated. This analysis extends the previous analysis of pairwise model combinations by expanding the set of ensembling methods and number of models included. Although results for multiple ensemble techniques were generated, only the method which included the combination with the lowest predicted RMSE is visualized here (weighting inversely proportionate to RMSE), with the rest being included in the supplementary materials.

### Identifying the Optimal Model Average Composition

As the optimal ensemble may not include all model types, differing the combinations of models, number of models, and weights used for averaging were explored. Specifically, inclusion of between one and eight model types, and 0-10 replicates of each (sampled from the available replicates with replacement), were used in a weighted average with weights being uniform, uniform with respect to the included model types, or inversely proportionate to prediction standard deviation, variance, or training set RMSE. To identify the optimal combination, a simple genetic algorithm was implemented to explore this parameter space to select combinations based on their expected RMSE.

In brief, this method mimics evolution to explore a high dimensional space by beginning with a “population” of parameter sets that undergo cycles of “selection” and “reproduction with mutation.” A starting population of 100 parameter sets were generated with uniform likelihood for all parameter values. The performance of each set was estimated by randomly drawing with replacement the prescribed number of model replicates for each model type, ensembling these predictions, calculating RMSE, and repeating this procedure to produce 50 estimates of RMSE.

The mean of this RMSE sample distribution was taken as the expected RMSE of the parameter set.

After evaluating the initial sets, 300 rounds of selection, reproduction with mutation, and evaluation were conducted. In each iteration 20 parameter sets were selected. Of these 15 corresponded to those with the lowest RMSE and 5 were drawn at random as a hedge against the algorithm becoming stuck in a local minimum. These parameter sets were mutated with at least one parameter being randomly altered. For each set the number of mutated parameters was drawn from a geometric distribution with p=0.5, meaning that most frequently one parameter would be changed but occasionally many would be altered. As with drawing 5 parameter sets at random this serves to increase the diversity of the parameter sets evaluated each cycle. After completion the top ten parameter sets were considered.

### Code availability

The analysis was conducted using Python (Van Rossum and Drake 2009) and relied on the use of common libraries for analysis (NumPy (Harris et al. 2020), scikit-learn (Pedregosa et al. 2011)) and initial visualizations (Plotly (Inc 2015)). Figures were produced using R (R Core Team 2021), benefiting from the drc (Ritz et al. 2015), Tidyverse (Wickham et al. 2019), ggrastr (Petukhov et al. 2021), and Patchwork (Pedersen 2020) libraries. Inkscape (inkscape.org) was used for final aesthetic adjustments. Environments for both languages were managed using Anaconda (2021). Scripts are available via Zenodo (DOI: 10.5281/zenodo.7697380).

## Results

### Pairwise ensembles often, but not always, outperform constituent models

Ensemble models composed of pairwise model averages generally outperformed their component models as assessed by average RMSE **(Figure 1 A)**. This was true for all models tested (LM (5/7 possible pairwise ensembles), BLUP (5/7), KNN (6/7), RNR (7/7), RF (7/7), SVR (7/7), DNN-CO (4/7), DNN-SO (6/7)). Further, the best performing ensemble models included the best performing constituent (or parent) models with the first, second, and fourth best performing ensemble models containing two of the top three individual models (BLUP, DNN- CO, LM). All ensembles that performed better than the best single model (BLUP) contained at least one of the top two models (BLUP or DNN-CO). While ensembling improved performance on average, some individual ensemble models had performance values intermediate to their component models, but no observed ensembles had an average performance worse than both of their constituent models.

**Figure 1.**
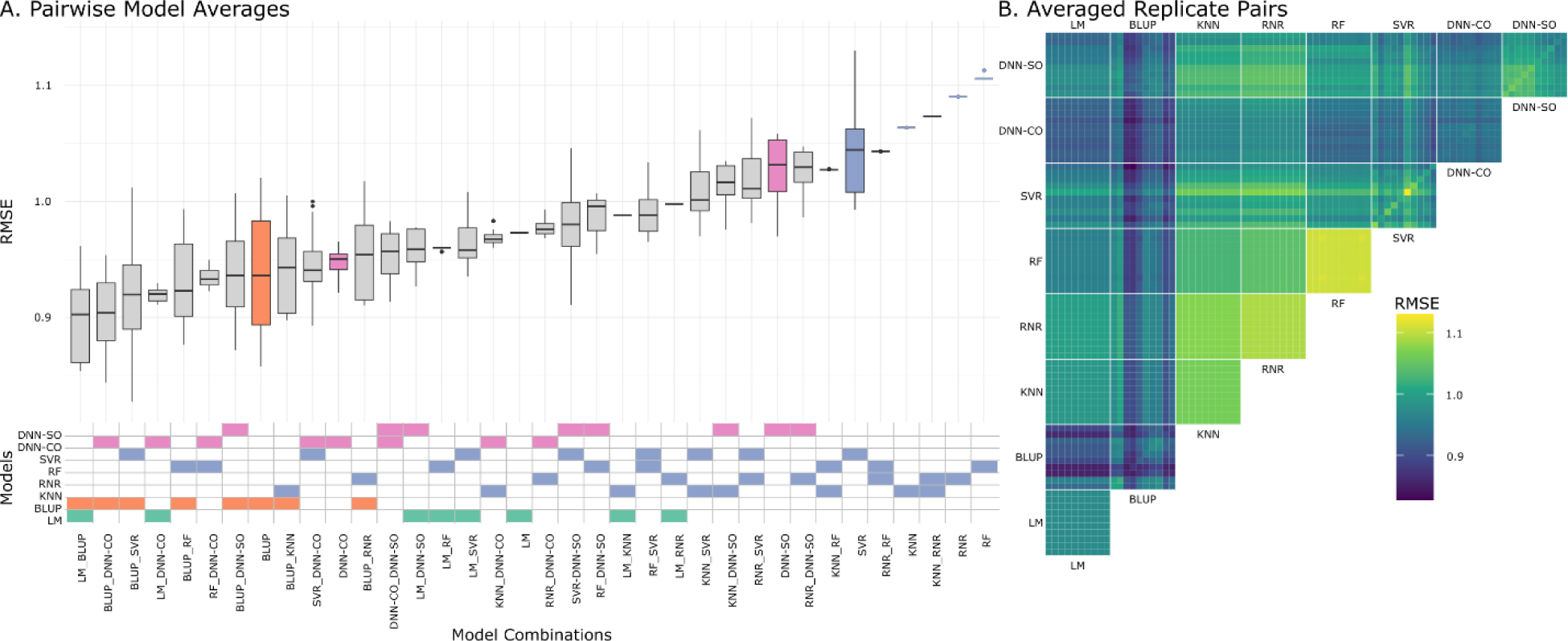
Performance of Pairwise Model Averages **A.** Performance of non-ensembled models of each type (filled boxplots) and pairwise combinations of model averages. Included models are written and shown graphically below. **B.** Pairwise combinations of all models and model replicates. Tile color indicates RMSE.

Except for a few cases, ensemble performance was variable depending on the specific model replicates (the same model run multiple times) included. With a few exceptions (LM, KNN, RNR) the tested models did not produce identical predictions across their replicates owing to models not always converging on the same set of parameters. This variability in performance within a model type resulted in variable performance when ensembling model types **(Figure 1 A, B)**. As a result, specific replicates of a model sometimes outperformed the average for an ensemble containing that given model type. One might envision a model replicate that outperforms not only the average of an ensemble but of *all* ensembles containing it. No instances of this were observed, i.e., where a specific model replicate had lower error than all ensembles that include it (**Figure 1 B** diagonal RMSE tiles vs off diagonal RMSE tiles). In the closest instance to this (BLUP replicate 2), only 14 ensembles (17.7%) outperformed the base model. The converse scenario, where a given model is outperformed by all ensembles which include it was much more common. Fourteen models (BLUP replicates 1, 6, 7, SVR replicates 0, 2, 3, 5, 6, 9, and DNN-SO replicates 0, 1,2, 3, 4) were improved by ensembling with *any* other model replicate (**Supplementary Table 1**). This indicates that the mean performance of a model type is not necessarily indicative of whether a model replicate is expected to be improved by ensembling as the model with the best average performance (BLUP) contained replicates which could be improved by ensembling with any other model.

Overall, model predictions had a 76.756% chance of being improved through a pairwise ensemble. The chance of improvement varied substantially by model type, but roughly increased as a function of average RMSE. The top three most performant model types, all had a greater than 50% chance of being improved: BLUP 63.418%, DNN-CO 57.848%, LM 53.165%, with the less performant models having a likelihood of being improved greater than 75% (DNN- SO 87.089%, SVR 96.962%, KNN 75.949%, RNR 88.608%, RF 91.013%) (**Table 1**). In summary, if a researcher or breeder randomly selected two of the included models for ensembling it would result in more accurate predictions than the two selected models on their own 76.756% of the time (within the models and datasets here tested). If a high performing model were used initially, adding a second model would still be expected to increase performance, albeit with a lower likelihood.

**Table 1.** Pairwise Model Averaging Results in More Performant Models with >50% Likelihood Likelihood of a replicate of a given model type being improved by ensembling with another model (including replicates of the same type) using a model average. The same as the average number of ensembles (out of 79 combinations) expected to be more performant. Ordered by ascending percentage.

| Model Type | Percent of Ensembles More Performant | Expected More Performant Ensembles (x/79) |
| --- | --- | --- |
| LM | 0.531646 | 42 |
| DNN-CO | 0.578481 | 45.7 |
| BLUP | 0.634177 | 50.1 |
| KNN | 0.759494 | 60 |
| DNN-SO | 0.870886 | 68.8 |
| RNR | 0.886076 | 70 |
| RF | 0.910127 | 71.9 |
| SVR | 0.96962 | 76.6 |

### Increasing the Number of Ensembled Models Often Improves Ensemble Performance

Various combinations of more than two models (or model replicates) together were tested to determine the utility of multi-model ensembles with different numbers of models and model types. Increasing the number of models or model replicates used, without consideration for the model type resulted in either no meaningful change in error (**Figure 2 A.**) or a lower expected error of the ensemble on average as more models were added (**Supplementary Figures 1-4 A.**), with the weighting scheme altering the trend. Additionally, the variability of ensembles errors’ is decreased with additional models. Where improvements in error were seen, the improvements associated with adding additional models diminished as the number of models increased. The improvement differed based on ensembling technique with the exponential decay’s “e” term (here representing the number of models required to reach ∼63% of the asymptote) varying from as high as 8876.604 and as low as 1.979 when using weights inversely proportionate to RMSE and using equal weights for each model type respectively (**Supplementary Table 2**.). This indicates that in practice there is limited utility in combining more than 6 randomly chosen models as all weighting schemes will have achieved >90% of the expected benefit except for the scheme weighting inversely proportionate to RMSE which showed little improvement with additional models.

**Figure 2.**
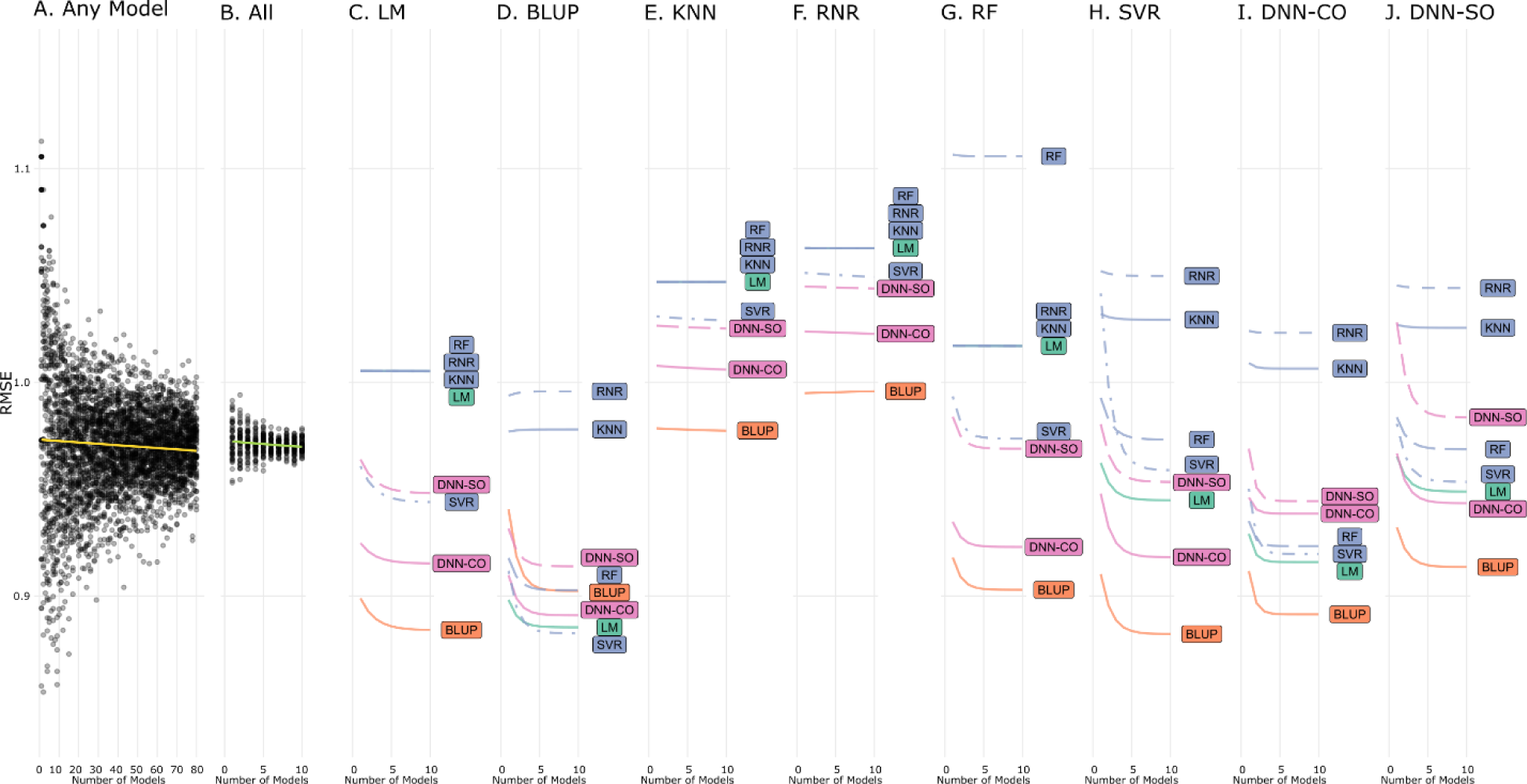
Performance of Ensembles by Number and Type of Models Used and Weighted Inversely Proportionate to RMSE. **A.** Performance from ensembles of model replicates selected at random from all trained models. Ensembling was performed using a weighted average with weights inversely proportionate to training set RMSE. Performance of 50 independent ensembles were simulated per step in number of models used. **B.** Performance of ensembles using 1-10 replicates of each model type. Note that the x axis here refers to the number of models in of an individual type – thus at x=10, 80 models are required. **C.-J.** Performance of ensembles using no more than two model types. Combinations which exhibited saturation with an increase in the number of models used were fit with an exponential decay function while the rest were fit as a linear regression. Note that the x axis here refers to the number of models in of an individual type – thus at x=10, 20 models are required.

When an equal number of replicates across all model types were ensembled, a similar trend to that above was observed, albeit with far less variability (**Figure 2 B., Supplementary Figures 1-4 B.**). A complementary analysis was done using all models trained and being different from the visualized ensembles in that sampling was not done with replacement.

Ensembles with equal model representation were considered using the weighting schemes previously described and using a multilayer perceptron to aggregate predictions (**Table 2**). Of the schemes examined, weighting replicates inversely proportional to each replicates’ variance resulted in the lowest error (0.923 RMSE). This represented a decrease in error of 0.047 RMSE relative to the average error of the best performing model type (BLUP, 0.937 RMSE, see (Kick et al. 2023)). Setting weights inversely proportionate to a replicate’s standard deviation was found to be less effective (0.925 RMSE) followed by uniform weighting (0.929 RMSE).

**Table 2.** Performance of Ensembling Strategies Performance for predictions from the training set and test set using all models and replicates is shown along with the percent RMSE relative to the average RMSE of the best single model (BLUP). The lowest error in each column is bolded. Predictions are weighted uniformly (Uniform), inversely proportionate to training set error (α RMSE-1), inversely proportionate to the standard deviation across model replicates (down weighting more similar models) (α s-1), and using a simple multilayer perceptron (MLP).

| Weighting | Train RMSE | Test RMSE | Percent Best Expected RMSE |
| --- | --- | --- | --- |
| Uniform | 0.501511 | 0.929173 | 0.991238 |
| $\alpha 1/s$ | 0.483222 | 0.925458 | 0.987275 |
| $\alpha 1/s^2$ | 0.513586 | **0.922683** | **0.984315** |
| $\alpha 1/RMSE$ | 0.319675 | 0.970065 | 1.034861 |
| MLP | **0.196820** | 1.062491 | 1.133461 |

Weighting inversely proportionate to a model’s training set error or aggregating predictions using a multilayer perceptron (1.062 RMSE) underperformed the expected RMSE for a non-ensemble BLUP model (0.937 RMSE). While weighting the contributions of different models to the ensemble showed potential for increasing accuracy, when all model types were used the potential improvements were modest. Given the variable performance across base models, and that certain model replicates are identical or effectively so, it is possible that a select subset of model types, or a variable number of each of these model types, would be more performant.

When combinations of specific model types were considered, variable performance gains were observed with respect to model combination and number (**Figure 2 C.-J.**). Many combinations outperform random ensembles (**Figure 2 A.**) and balanced ensembles (**Figure 2 B.**) – and do so with fewer model replicates included in the ensemble. Combinations shown in panels **C.-J.** contain at most 20 models (i.e., up to 10 of each for each pair) whereas panels **A.** and **B.** contain up to 80. Similar trends are seen across ensembling methods (**Supplementary Figures 1-4**), but performance varies. The combination with the lowest asymptotic error (“c” term) (BLUP & SVR) used inverse RMSE weighting (**Figure 2**), but the average asymptotic error was *greatest* for ensembles in this group (0.978 RMSE), while all other methods had an average of 0.966 RMSE. Inverse RMSE weighting appears to produce more variable results with a standard deviation across “c” terms of 0.058 RMSE, whereas assigning weights uniformly (0.049 RMSE), uniformly with respect to model type (0.049 RMSE), or inversely proportionate to standard deviation (0.046 RMSE), or variance (0.046 RMSE) all had lower standard deviations.

As with the results for pairwise ensembles, certain model types performed best when combined with other model types (**Figure 2 C.-J.**). For example, most model types were improved to the greatest extent when combined with BLUP, DNN-CO, or LM models. These model types were some of the most accurate models when tested on their own, so that may at least partially explain their utility in ensemble models.

Expanding on the above, ensembles with unequal representation of between one and eight model types were considered and evaluated. All non-multilayer perceptron ensembling strategies were considered. Because exploring all possible combinations of models and weighting schemes is infeasible an optimization approach was used instead. A genetic algorithm was used to select the number of replicates of each model (selected at random with replacement) and the weighting used to combine them. The best ensemble identified resulted in an RMSE of 0.872 and was composed of 2 linear models, 8 BLUPs, 4 random forests, 2 support vector regressions and 6 DNN-CO deep neural networks averaged with weights inversely proportionate to training set RMSE (**Supplementary Table 3**). This represents an improvement of -0.010 RMSE relative to the best ensemble of two model types (BLUP and SVR, 0.882 RMSE), and -0.098 RMSE relative to the best model without ensembling (BLUP, 0.937 RMSE). In percentages, using the genetic algorithm optimized ensemble resulted in a 1.134% lower error relative to the best two model ensemble and 6.980% lower error relative to the best non-ensembled model.

Among the ten most optimal ensembles, all used weights inversely proportionate to the RMSE, similar to what was seen in the performance of this approach in ensembles of two models. Ensemble composition, although more variable among top ensembles, appears to be key. None of the top ensembles included the model types absent from the best combination (KNN, RNR, DNN-SO). Thus, model type, number of models, and ensembling strategy all contribute to ensemble performance.

## Discussion

### Ensembling Improves Yield Predictions

Combining predictions from genetic, environment, and management containing linear, best linear unbiased predictor, deep neural network, and machine learning models frequently resulted in better accuracy than the base models. This effect appears under many variations on ensembles: using multiple models, either of differing model types or the same, and applying differing weighting schemes. While there appear to be more and less optimal ensembling methods, ensembling generally outperforms the performance of base models. Based on our results, a researcher or breeder would, more often than not, be better off choosing two models at random and ensembling them together than using any one model on its own. That said, it is important to note that these results may not hold true in all datasets and our conclusions are confined to scenarios were environmental, as well as genetic information, is explicitly modeled and datasets where previous studies have shown that environmental variation is extremely important.

While it is common in evaluating phenotypic prediction models to benchmark a model of interest against other models, it is less common to consider the improved accuracy which may be achieved through using many or all of the trained models.. Focusing on finding the single *“best”* model results in missing out on readily available predictive power. Importantly these ensembling approaches require no major modifications to experimental design or analysis, being easily added to an existing workflow if that workflow already contains multiple modeling types, or replication of the same model type with different parameters. Moving from selecting a single “*best*” to instead selecting the “*best sets*” of models for a given application enables individual models to be optimized to represent specific features in the data rather than requiring one framework to excel at modeling the full complexity of the problem.

While model ensembling benefits from incorporation of models that capture different aspects of the data, this is not a requirement for a performance increase. These results show that when the same model has variability between replicates (replicates do not converge to the same or effectively the same parameter values) then merely ensembling replicates is sufficient to elicit a performance increase. While it is possible to create an ensemble using a single model type, and most of the weighting schemes tested outperformed the base models, optimal results require a diversity of model types. Combining replicates from the top three performing models BLUP, DNN-CO, LM with lower performing SVR and RF models resulted in the best ensemble. This is a diverse set of modeling frameworks, and the diversity of models likely contributes to the performance – if a method’s performance is poor in a certain subset of the data it is buffered by predictions from methods which are unlikely to struggle on the exact same subset.

### Designing Experiments for Ensembling

The best performing ensembling approaches tested may all be used without modification to standard data partitioning approaches (e.g., separating the data into a training and testing set or using multiple cross validation folds). Modification of these approaches to include a validation set (or an additional validation set if hyperparameter tuning is being conducted) for ensemble selection may further improve ensemble performance. Combining predictions with a weighted average proportionate to inverse RMSE or the use of a multilayer perceptron are methods which may be undermined if base models are overfit. In that case the most influential models would be those *least likely to generalize* to new observations leading to a higher out of sample error. If model performance on a validation set were used instead then this issue would be avoided, and the performance of these methods would be expected to increase.

Beyond designing data partitioning with ensembling in mind, training models to operate on the output of other models (“model stacking”) has the potential to improve performance, but also implicate cases in which individual models or modeling frameworks are of greater use. Key covariates could be provided to these models in addition to providing predicted values. By so doing, models which perform especially well or especially poorly in specific cases (e.g., certain climates, cultivars, management treatments, etc.) could be identified by examining the ensemble model. Beyond improving performance this approach would provide nuance to model recommendations; rather than identifying the model with the best overall performance, the best model for specific locations or uses could be determined.

Due to the ease of implementation and clear performance benefits we recommend the further study of ensembling methods for predictive problems in agriculture broadly and for phenotypic prediction specifically. Further investigation of the combinations of modeling frameworks which best complement each other, aggregation methods that confer greater accuracy and robustness, and data partitioning strategies which balance ensemble selection with available data for model training, are expected to confer further benefits in diverse fields and applications.

## Statements and Declarations

### Author Contributions

The computational study was designed by DRK with input from JDW. The manuscript was written by DRK and edited by JDW and DRK.

### Data Availability Statement

Data for maize phenotypes and genomic, soil, weather, and management covariates came from the Genomes to Fields Initiative data (McFarland, et al. 2020). These data are publicly available in their raw form through the CyVerse Discovery environment and a cleaned version of these data are available through Zenodo (zenodo.org/record/6916775 DOI 10.5281/zenodo.6916775). Model predictions were made using the models document in Kick et al. 2023 (Kick et al. 2023) and are available along with the analysis scripts written for the current study through Zenodo (DOI: 10.5281/zenodo.7697380).

### Ethics approval

This study did not involve human or animal subjects and thus was not reviewed by an institutional or national research ethics committee.

### Consent to participate

This study did not involve participants and thus informed consent of the research subjects was not collected.

### Consent to publish

The authors affirm that human research participants were not involved and thus no consent to publish was collected.

## Key Message

Predicting a phenotype from aggregating multiple models can be more accurate than a single model. Using more model types, replicates, and weighting models inversely proportionate to error is beneficial.

### Acknowledgements

This research used resources provided by the United States Department of Agriculture’s Agricultural Research Service (project number 5070-21000-041-000-D). The SCINet project of the USDA Agricultural Research Service (project number 0500-00093-001-00-D) was instrumental in the training of the models used in this work. In addition, we would like to acknowledge those presently and historically involved in generating data for the Genomes to Fields Initiative.

---

**Supplementary Figure 1.**
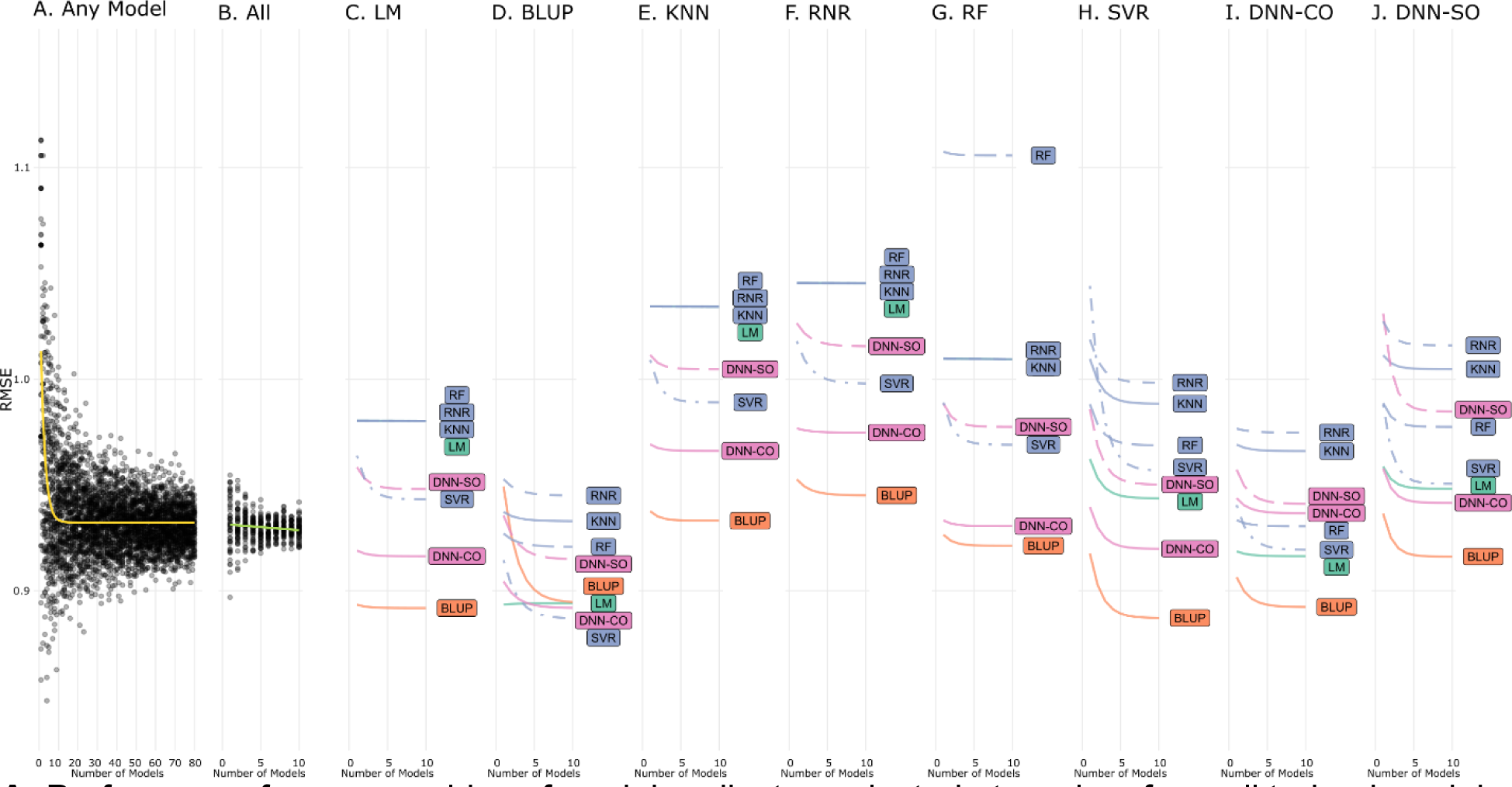
Performance of Ensembles by Number and Type of Models Used and Weighted Uniformly. **A.** Performance from ensembles of model replicates selected at random from all trained models. Ensembling was performed using a simple arithmetic average. Performance of 50 independent ensembles were simulated per step in number of models used. **B.** Performance of ensembles using 1-10 replicates of each model type. Note that the x axis here refers to the number of models in of an individual type – thus at x=10, 80 models are required. **C.-J.** Performance of ensembles using no more than two model types. Combinations which exhibited saturation with an increase in the number of models used were fit with an exponential decay function while the rest were fit as a linear regression. Note that the x axis here refers to the number of models in of an individual type – thus at x=10, 20 models are required.

**Supplementary Figure 2.**
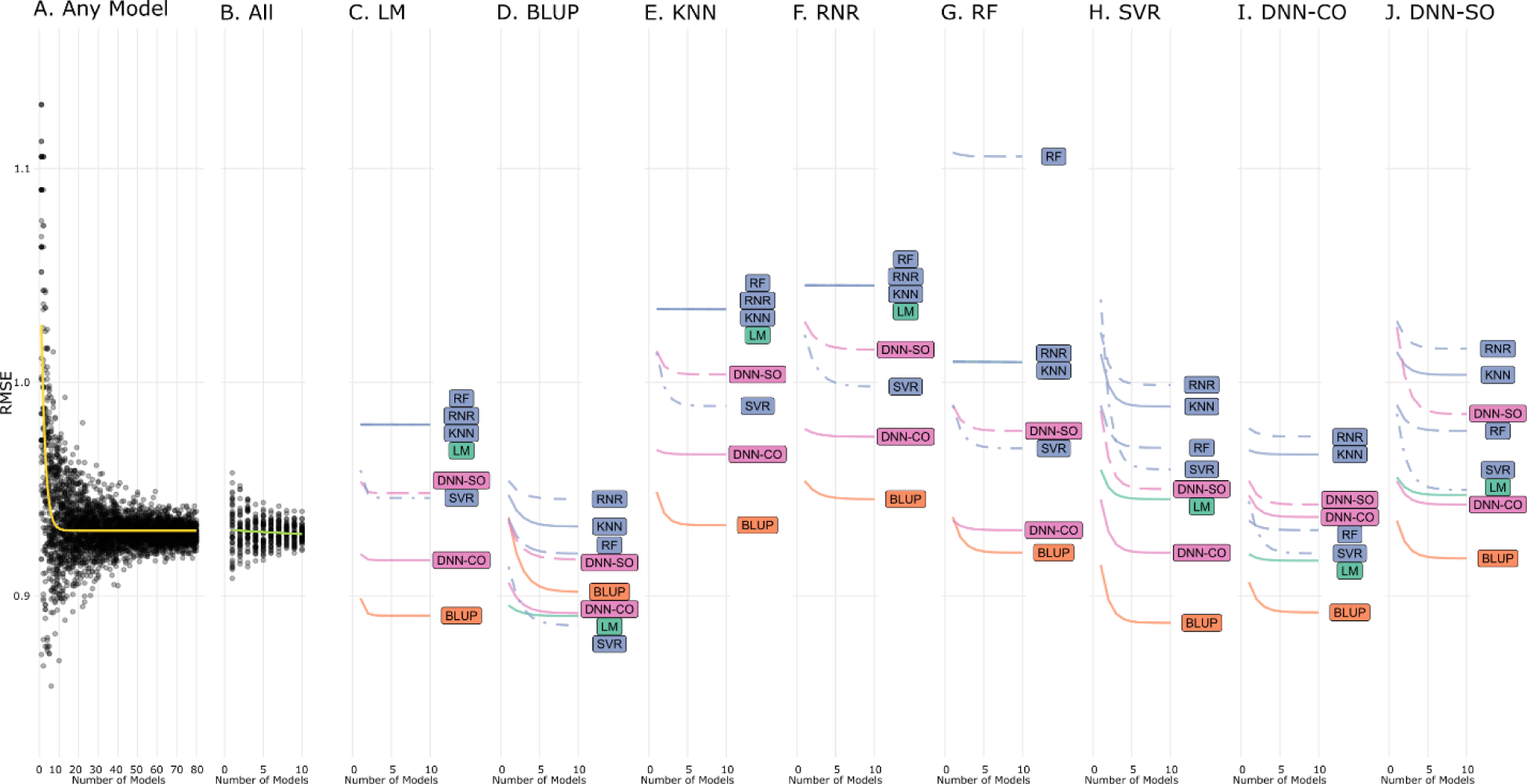
Performance of Ensembles by Number and Type of Models Used and Weighted Uniformly By Model Type. **A.** Performance from ensembles of model replicates selected at random from all trained models. Ensembling was performed using a weighted average with each model type receiving equal total weight. Performance of 50 independent ensembles were simulated per step in number of models used. **B.** Performance of ensembles using 1-10 replicates of each model type. Note that the x axis here refers to the number of models in of an individual type – thus at x=10, 80 models are required. **C.-J.** Performance of ensembles using no more than two model types. Combinations which exhibited saturation with an increase in the number of models used were fit with an exponential decay function while the rest were fit as a linear regression. Note that the x axis here refers to the number of models in of an individual type – thus at x=10, 20 models are required.

**Supplementary Figure 3.**
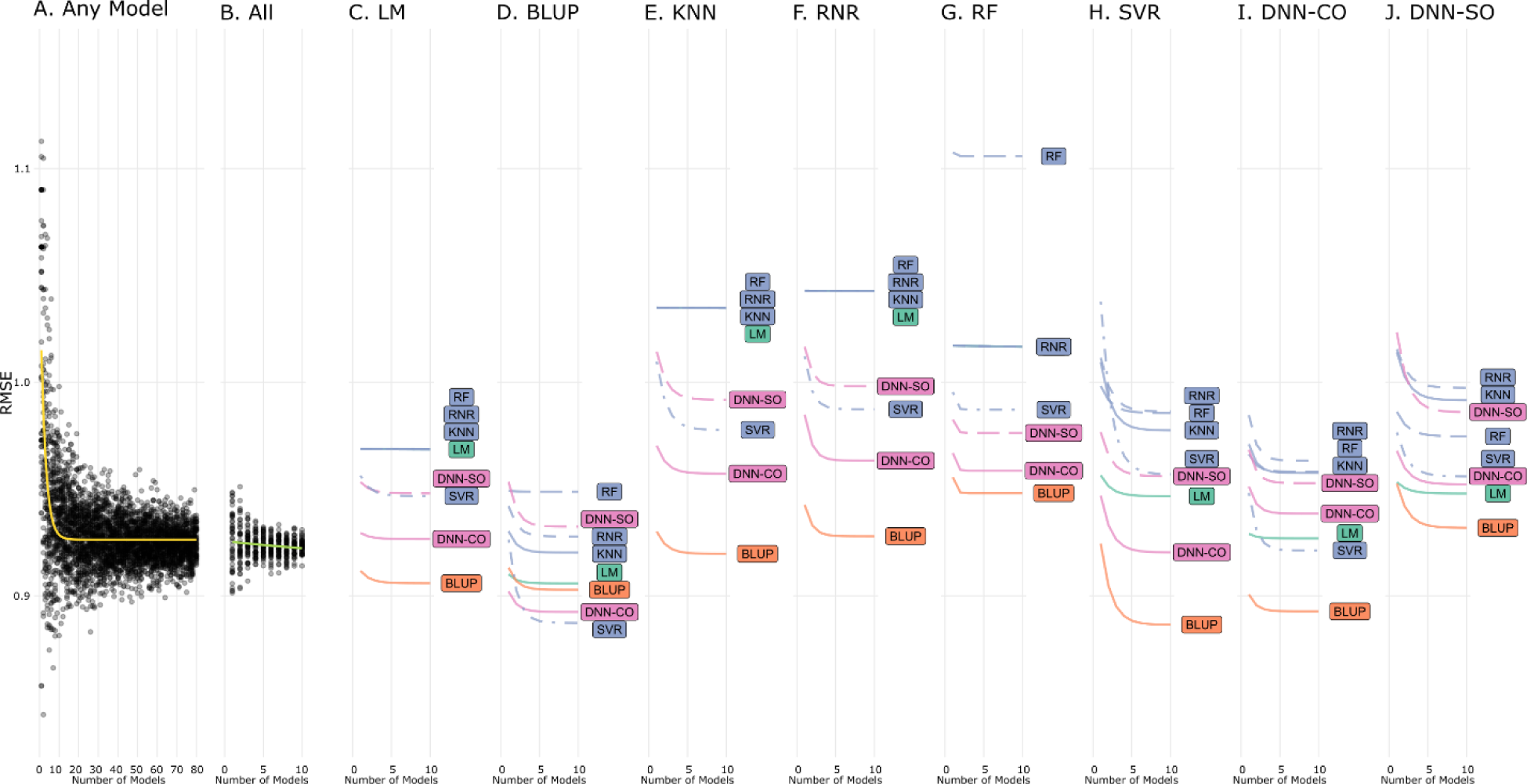
Performance of Ensembles by Number and Type of Models Used and Weighted Inversely Proportionate to Variance. **A.** Performance from ensembles of model replicates selected at random from all trained models. Ensembling was performed using a weighted average with weights inversely proportionate to the variance of predictions. Performance of 50 independent ensembles were simulated per step in number of models used. **B.** Performance of ensembles using 1-10 replicates of each model type. Note that the x axis here refers to the number of models in of an individual type – thus at x=10, 80 models are required. **C.-J.** Performance of ensembles using no more than two model types. Combinations which exhibited saturation with an increase in the number of models used were fit with an exponential decay function while the rest were fit as a linear regression. Note that the x axis here refers to the number of models in of an individual type – thus at x=10, 20 models are required.

**Supplementary Figure 4.**
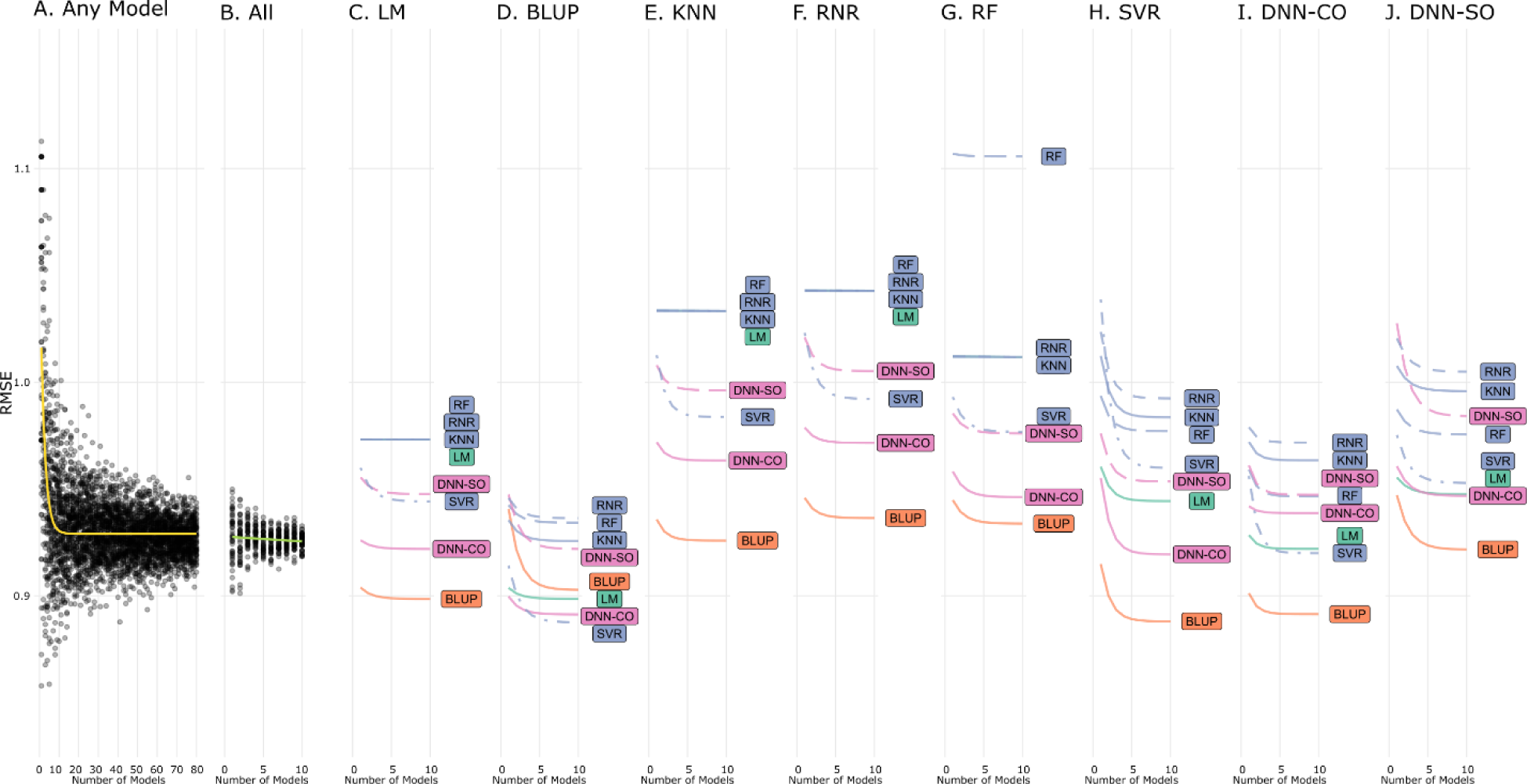
Performance of Ensembles by Number and Type of Models Used and Weighted Inversely Proportionate to Standard Deviation. **A.** Performance from ensembles of model replicates selected at random from all trained models. Ensembling was performed using a weighted average with weights inversely proportionate to the standard deviation of predictions. Performance of 50 independent ensembles were simulated per step in number of models used. **B.** Performance of ensembles using 1-10 replicates of each model type. Note that the x axis here refers to the number of models in of an individual type – thus at x=10, 80 models are required. **C.-J.** Performance of ensembles using no more than two model types. Combinations which exhibited saturation with an increase in the number of models used were fit with an exponential decay function while the rest were fit as a linear regression. Note that the x axis here refers to the number of models in of an individual type – thus at x=10, 20 models are required.

**Supplementary Table 1.**
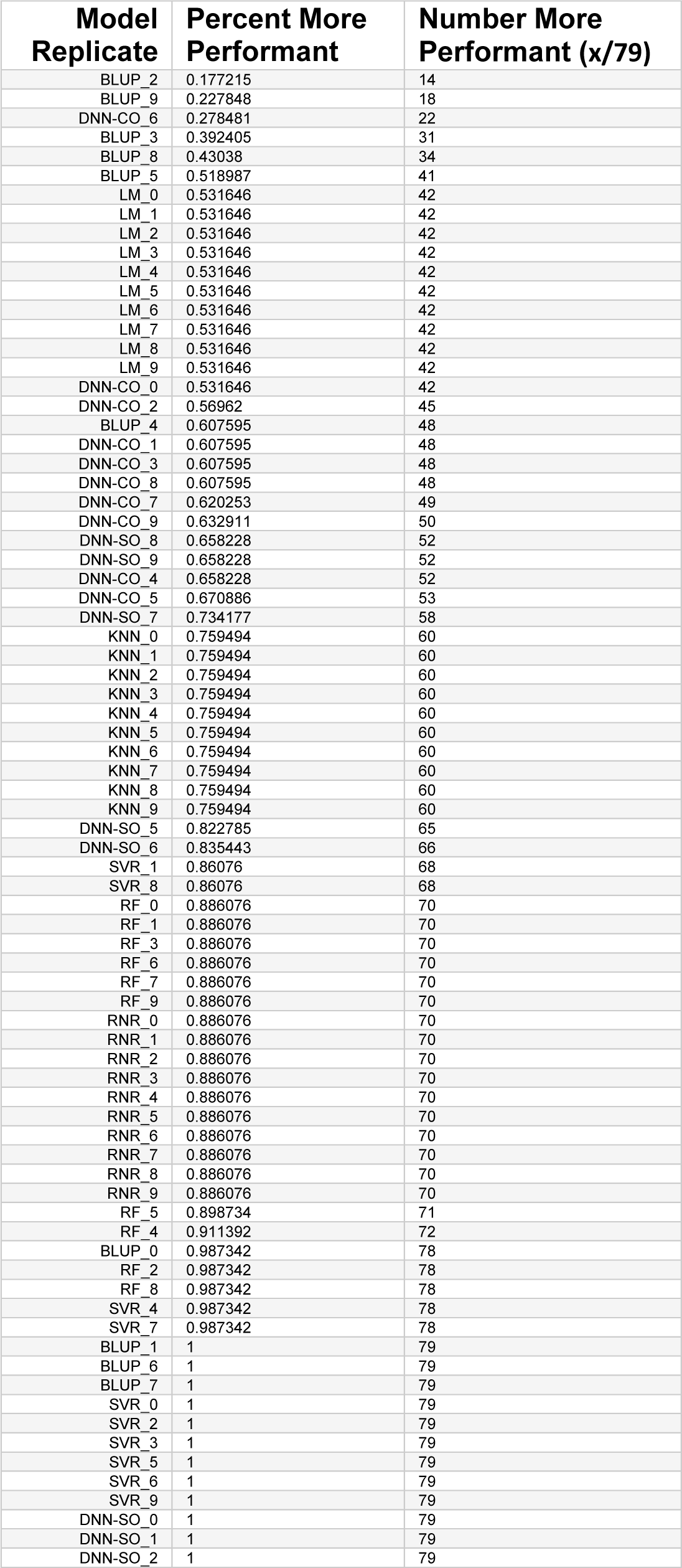
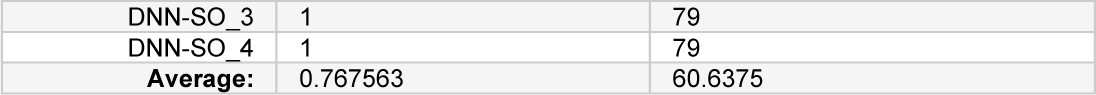
Model Replicate Performance Relative to Performance of Model Averages Performance of each model replicate with the percent and number of pairwise averages which resulted in a RMSE less than the base model. Sorted by the percentage of pairwise averages more performant.

**Supplementary Table 2.**
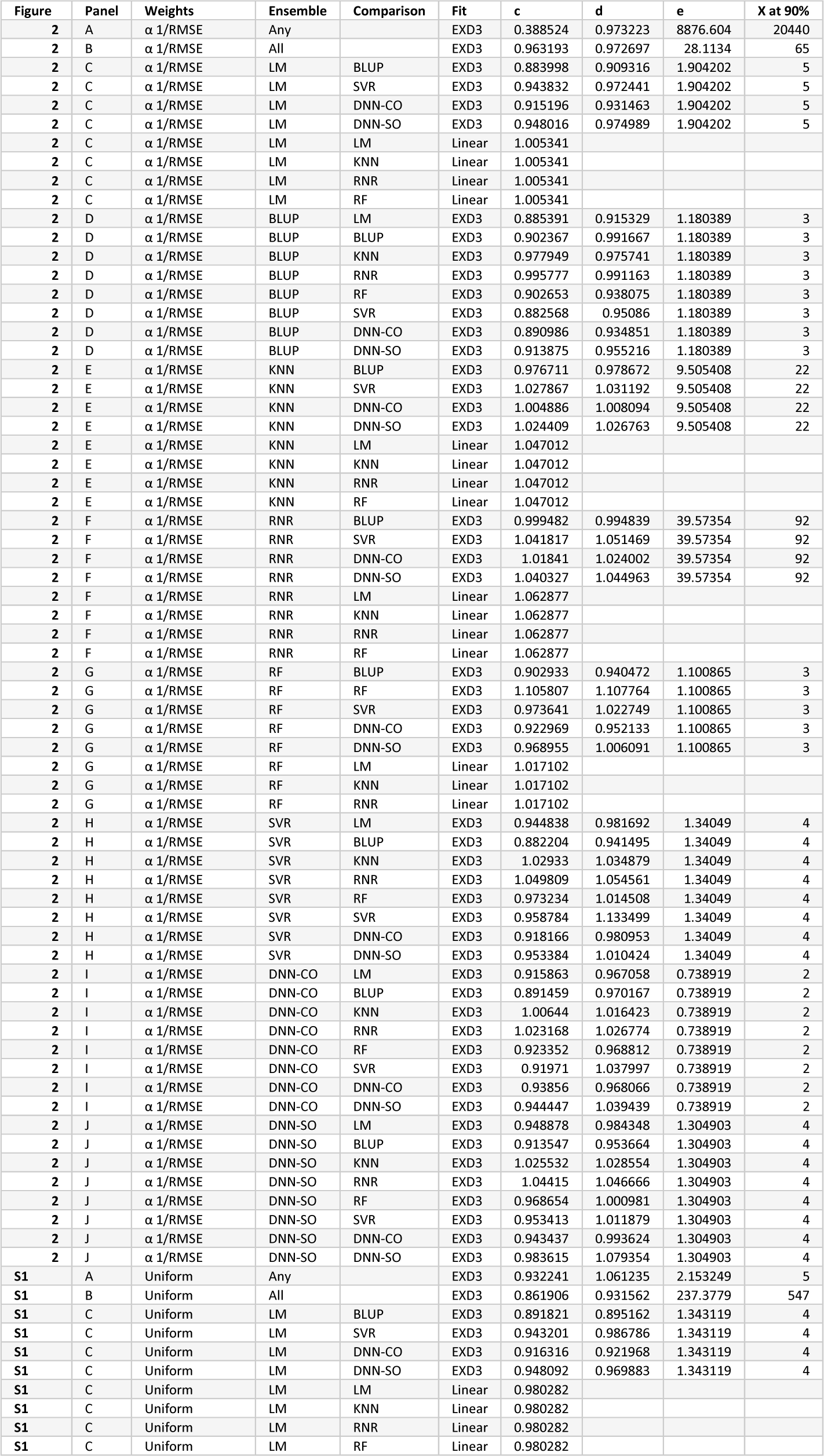
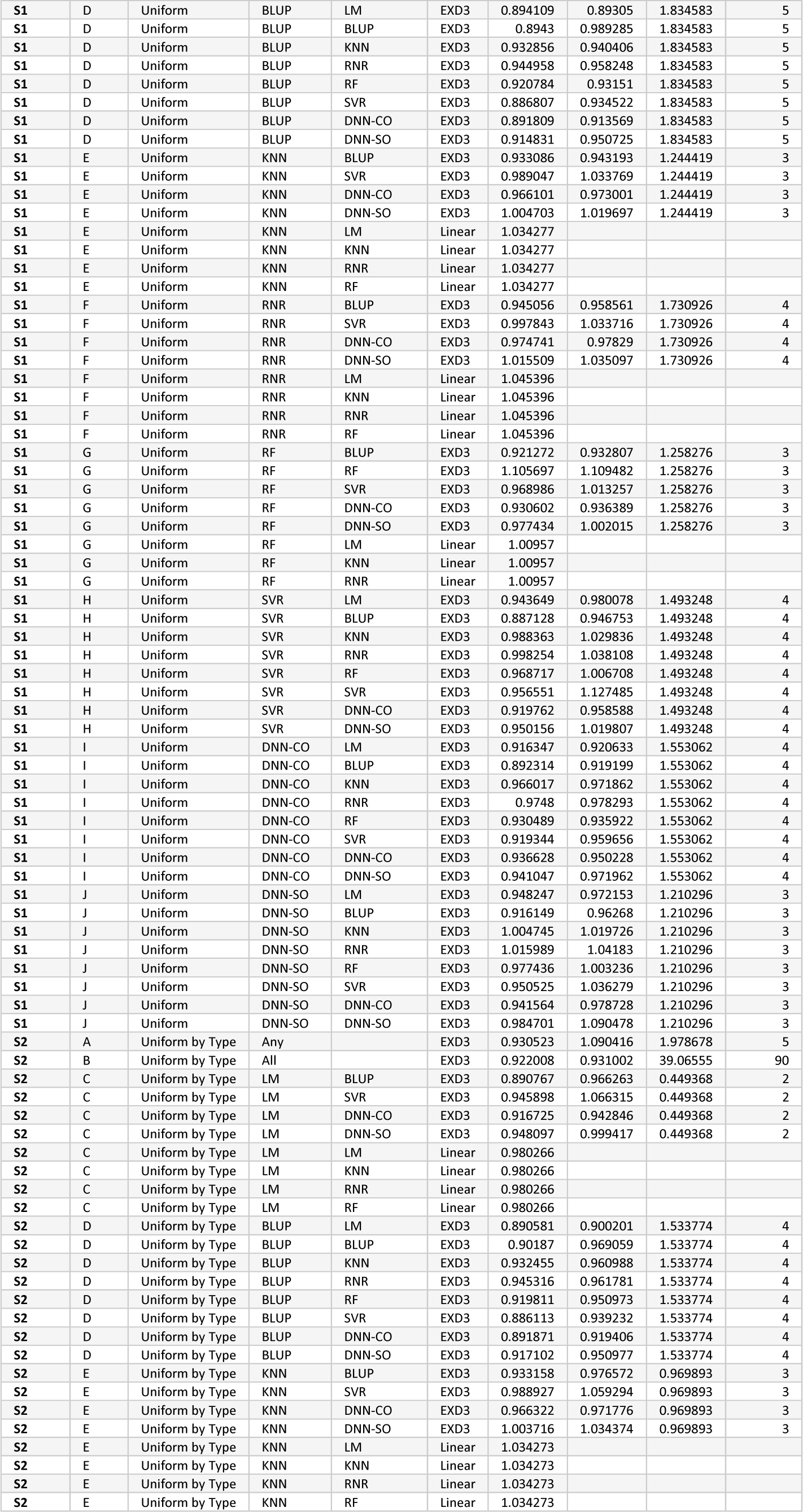
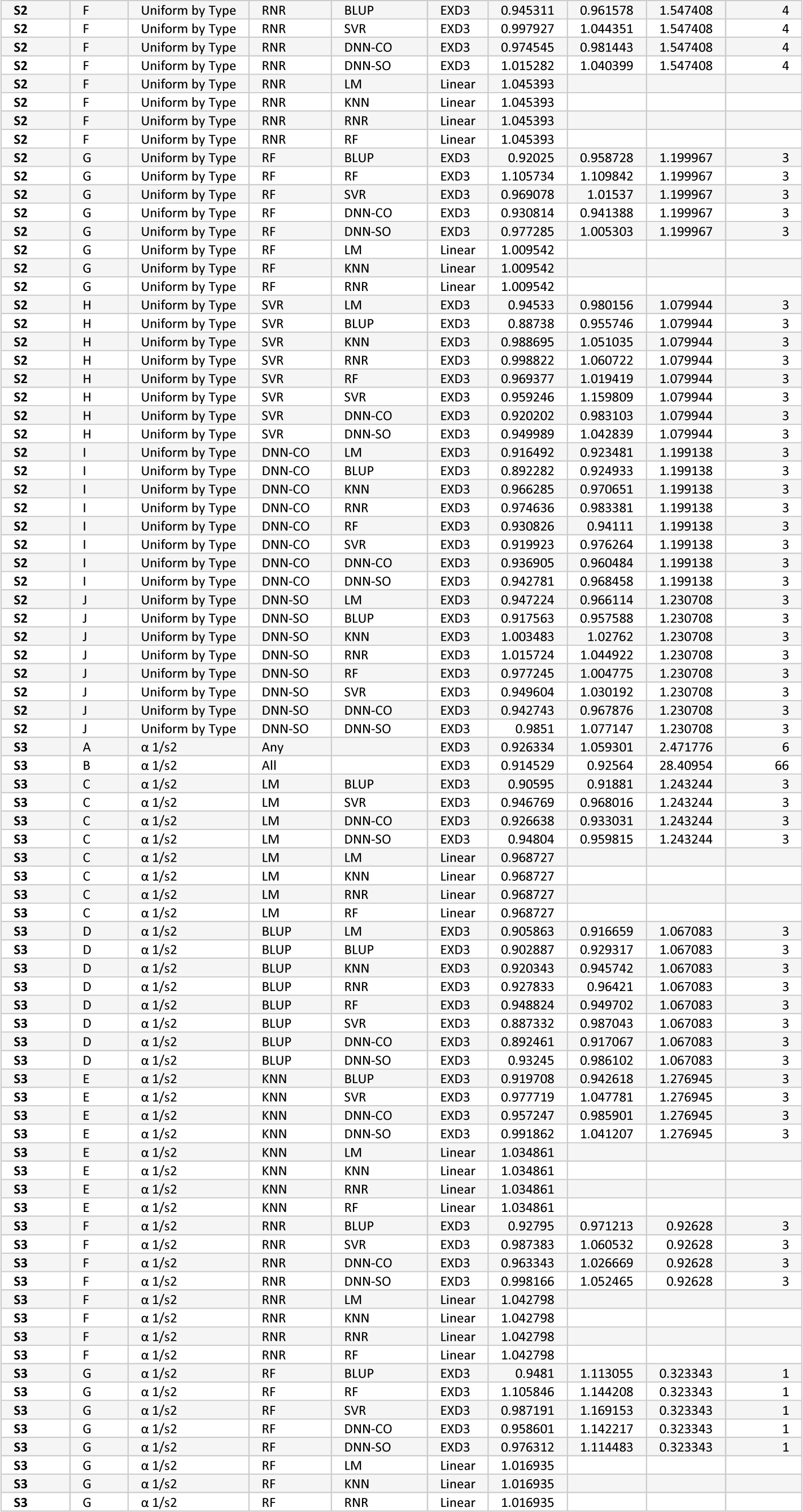
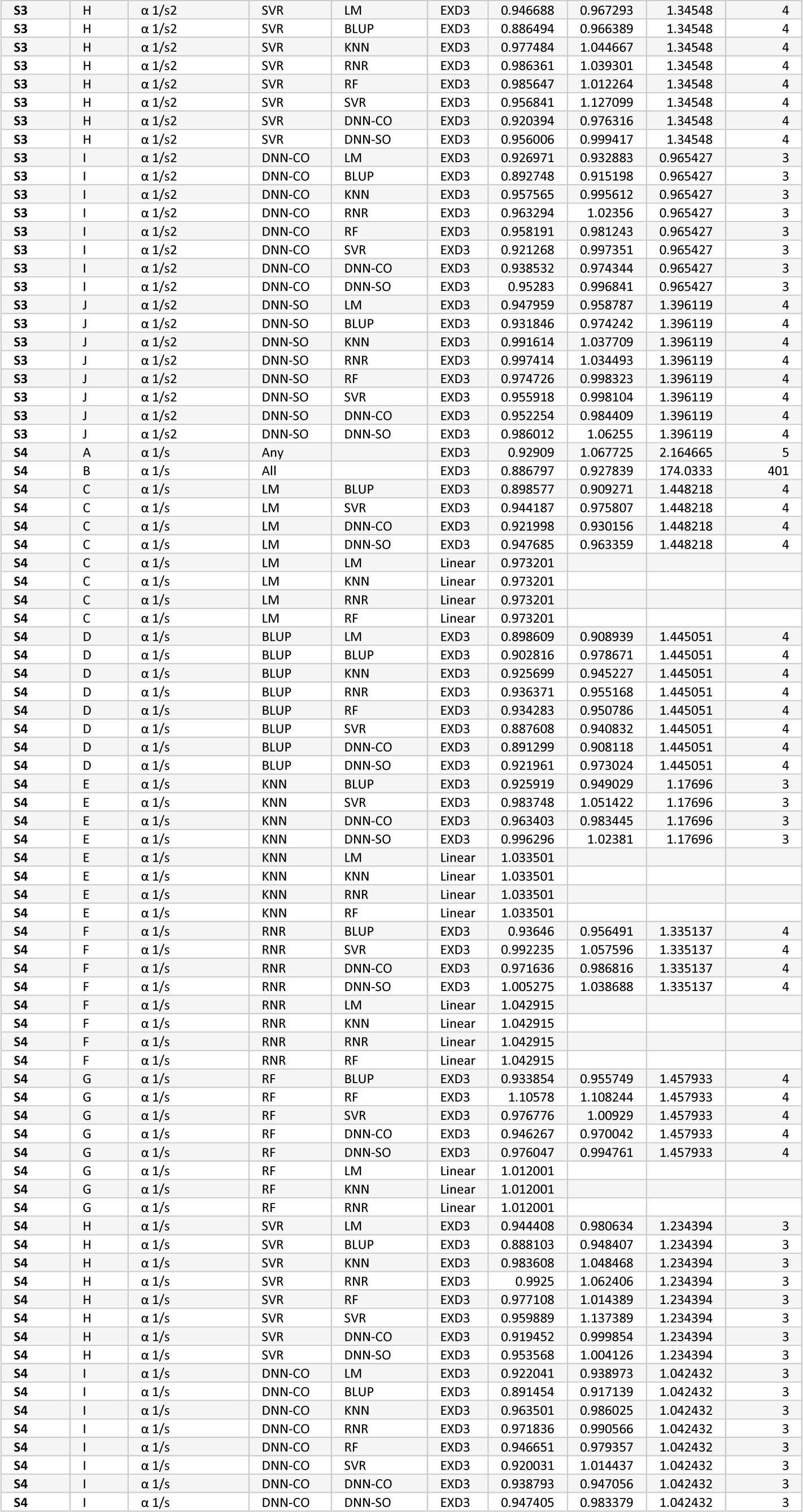
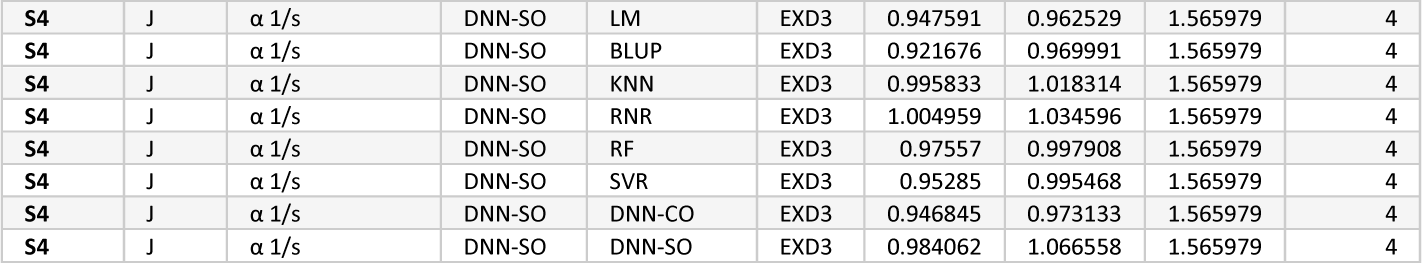
Ensembling Strategy Performance Trends Ensemble error was models as following an exponential decay or linear trend with respect to the number of models added into the ensemble. Linear fits were used if an ensemble’s variability was below a threshold. Parameters of the fits visualized in Figure 2 and Supplementary Figures 1-4 are shown. For convenience each row includes reference to the figure and panel that contains a visualization of that model. Intercept terms for linear models displayed in the “c” column. Additionally, the x axis value required to reach 90% of the asymptote is included. Note however that the number of models required for each is variable with x being equal to the number of individual models used when “Any” model type was included, 8x when “All” model types were included, and 2x when pairwise combinations of models were included.

| Figure | Panel | Weights | Ensemble | Comparison | Fit | c | d | e | X at 90% |
| --- | --- | --- | --- | --- | --- | --- | --- | --- | --- |
| 2 | A | $\alpha$ 1/RMSE | Any | | EXD3 | 0.388524 | 0.973223 | 8876.604 | 20440 |
| 2 | B | $\alpha$ 1/RMSE | All | | EXD3 | 0.963193 | 0.972697 | 28.1134 | 65 |
| 2 | C | $\alpha$ 1/RMSE | LM | BLUP | EXD3 | 0.883998 | 0.909316 | 1.904202 | 5 |
| 2 | C | $\alpha$ 1/RMSE | LM | SVR | EXD3 | 0.943832 | 0.972441 | 1.904202 | 5 |
| 2 | C | $\alpha$ 1/RMSE | LM | DNN-CO | EXD3 | 0.915196 | 0.931463 | 1.904202 | 5 |
| 2 | C | $\alpha$ 1/RMSE | LM | DNN-SO | EXD3 | 0.948016 | 0.974989 | 1.904202 | 5 |
| 2 | C | $\alpha$ 1/RMSE | LM | LM | Linear | 1.005341 | | | |
| 2 | C | $\alpha$ 1/RMSE | LM | KNN | Linear | 1.005341 | | | |
| 2 | C | $\alpha$ 1/RMSE | LM | RNR | Linear | 1.005341 | | | |
| 2 | C | $\alpha$ 1/RMSE | LM | RF | Linear | 1.005341 | | | |
| 2 | D | $\alpha$ 1/RMSE | BLUP | LM | EXD3 | 0.885391 | 0.915329 | 1.180389 | 3 |
| 2 | D | $\alpha$ 1/RMSE | BLUP | BLUP | EXD3 | 0.902367 | 0.991667 | 1.180389 | 3 |
| 2 | D | $\alpha$ 1/RMSE | BLUP | KNN | EXD3 | 0.977949 | 0.975741 | 1.180389 | 3 |
| 2 | D | $\alpha$ 1/RMSE | BLUP | RNR | EXD3 | 0.995777 | 0.991163 | 1.180389 | 3 |
| 2 | D | $\alpha$ 1/RMSE | BLUP | RF | EXD3 | 0.902653 | 0.938075 | 1.180389 | 3 |
| 2 | D | $\alpha$ 1/RMSE | BLUP | SVR | EXD3 | 0.882568 | 0.95086 | 1.180389 | 3 |
| 2 | D | $\alpha$ 1/RMSE | BLUP | DNN-CO | EXD3 | 0.890986 | 0.934851 | 1.180389 | 3 |
| 2 | D | $\alpha$ 1/RMSE | BLUP | DNN-SO | EXD3 | 0.913875 | 0.955216 | 1.180389 | 3 |
| 2 | E | $\alpha$ 1/RMSE | KNN | BLUP | EXD3 | 0.976711 | 0.978672 | 9.505408 | 22 |
| 2 | E | $\alpha$ 1/RMSE | KNN | SVR | EXD3 | 1.027867 | 1.031192 | 9.505408 | 22 |
| 2 | E | $\alpha$ 1/RMSE | KNN | DNN-CO | EXD3 | 1.004886 | 1.008094 | 9.505408 | 22 |
| 2 | E | $\alpha$ 1/RMSE | KNN | DNN-SO | EXD3 | 1.024409 | 1.026763 | 9.505408 | 22 |
| 2 | E | $\alpha$ 1/RMSE | KNN | LM | Linear | 1.047012 | | | |
| 2 | E | $\alpha$ 1/RMSE | KNN | KNN | Linear | 1.047012 | | | |
| 2 | E | $\alpha$ 1/RMSE | KNN | RNR | Linear | 1.047012 | | | |
| 2 | E | $\alpha$ 1/RMSE | KNN | RF | Linear | 1.047012 | | | |
| 2 | F | $\alpha$ 1/RMSE | RNR | BLUP | EXD3 | 0.999482 | 0.994839 | 39.57354 | 92 |
| 2 | F | $\alpha$ 1/RMSE | RNR | SVR | EXD3 | 1.041817 | 1.051469 | 39.57354 | 92 |
| 2 | F | $\alpha$ 1/RMSE | RNR | DNN-CO | EXD3 | 1.01841 | 1.024002 | 39.57354 | 92 |
| 2 | F | $\alpha$ 1/RMSE | RNR | DNN-SO | EXD3 | 1.040327 | 1.044963 | 39.57354 | 92 |
| 2 | F | $\alpha$ 1/RMSE | RNR | LM | Linear | 1.062877 | | | |
| 2 | F | $\alpha$ 1/RMSE | RNR | KNN | Linear | 1.062877 | | | |
| 2 | F | $\alpha$ 1/RMSE | RNR | RNR | Linear | 1.062877 | | | |
| 2 | F | $\alpha$ 1/RMSE | RNR | RF | Linear | 1.062877 | | | |
| 2 | G | $\alpha$ 1/RMSE | RF | BLUP | EXD3 | 0.902933 | 0.940472 | 1.100865 | 3 |
| 2 | G | $\alpha$ 1/RMSE | RF | RF | EXD3 | 1.105807 | 1.107764 | 1.100865 | 3 |
| 2 | G | $\alpha$ 1/RMSE | RF | SVR | EXD3 | 0.973641 | 1.022749 | 1.100865 | 3 |
| 2 | G | $\alpha$ 1/RMSE | RF | DNN-CO | EXD3 | 0.922969 | 0.952133 | 1.100865 | 3 |
| 2 | G | $\alpha$ 1/RMSE | RF | DNN-SO | EXD3 | 0.968955 | 1.006091 | 1.100865 | 3 |
| 2 | G | $\alpha$ 1/RMSE | RF | LM | Linear | 1.017102 | | | |
| 2 | G | $\alpha$ 1/RMSE | RF | KNN | Linear | 1.017102 | | | |
| 2 | G | $\alpha$ 1/RMSE | RF | RNR | Linear | 1.017102 | | | |
| 2 | H | $\alpha$ 1/RMSE | SVR | LM | EXD3 | 0.944838 | 0.981692 | 1.34049 | 4 |
| 2 | H | $\alpha$ 1/RMSE | SVR | BLUP | EXD3 | 0.882204 | 0.941495 | 1.34049 | 4 |
| 2 | H | $\alpha$ 1/RMSE | SVR | KNN | EXD3 | 1.02933 | 1.034879 | 1.34049 | 4 |
| 2 | H | $\alpha$ 1/RMSE | SVR | RNR | EXD3 | 1.049809 | 1.054561 | 1.34049 | 4 |
| 2 | H | $\alpha$ 1/RMSE | SVR | RF | EXD3 | 0.973234 | 1.014508 | 1.34049 | 4 |
| 2 | H | $\alpha$ 1/RMSE | SVR | SVR | EXD3 | 0.958784 | 1.133499 | 1.34049 | 4 |
| 2 | H | $\alpha$ 1/RMSE | SVR | DNN-CO | EXD3 | 0.918166 | 0.980953 | 1.34049 | 4 |
| 2 | H | $\alpha$ 1/RMSE | SVR | DNN-SO | EXD3 | 0.953384 | 1.010424 | 1.34049 | 4 |
| 2 | I | $\alpha$ 1/RMSE | DNN-CO | LM | EXD3 | 0.915863 | 0.967058 | 0.738919 | 2 |
| 2 | I | $\alpha$ 1/RMSE | DNN-CO | BLUP | EXD3 | 0.891459 | 0.970167 | 0.738919 | 2 |
| 2 | I | $\alpha$ 1/RMSE | DNN-CO | KNN | EXD3 | 1.00644 | 1.016423 | 0.738919 | 2 |
| 2 | I | $\alpha$ 1/RMSE | DNN-CO | RNR | EXD3 | 1.023168 | 1.026774 | 0.738919 | 2 |
| 2 | I | $\alpha$ 1/RMSE | DNN-CO | RF | EXD3 | 0.923352 | 0.968812 | 0.738919 | 2 |
| 2 | I | $\alpha$ 1/RMSE | DNN-CO | SVR | EXD3 | 0.91971 | 1.037997 | 0.738919 | 2 |
| 2 | I | $\alpha$ 1/RMSE | DNN-CO | DNN-CO | EXD3 | 0.93856 | 0.968066 | 0.738919 | 2 |
| 2 | I | $\alpha$ 1/RMSE | DNN-CO | DNN-SO | EXD3 | 0.944447 | 1.039439 | 0.738919 | 2 |
| 2 | J | $\alpha$ 1/RMSE | DNN-SO | LM | EXD3 | 0.948878 | 0.984348 | 1.304903 | 4 |
| 2 | J | $\alpha$ 1/RMSE | DNN-SO | BLUP | EXD3 | 0.913547 | 0.953664 | 1.304903 | 4 |
| 2 | J | $\alpha$ 1/RMSE | DNN-SO | KNN | EXD3 | 1.025532 | 1.028554 | 1.304903 | 4 |
| 2 | J | $\alpha$ 1/RMSE | DNN-SO | RNR | EXD3 | 1.04415 | 1.046666 | 1.304903 | 4 |
| 2 | J | $\alpha$ 1/RMSE | DNN-SO | RF | EXD3 | 0.968654 | 1.000981 | 1.304903 | 4 |
| 2 | J | $\alpha$ 1/RMSE | DNN-SO | SVR | EXD3 | 0.953413 | 1.011879 | 1.304903 | 4 |
| 2 | J | $\alpha$ 1/RMSE | DNN-SO | DNN-CO | EXD3 | 0.943437 | 0.993624 | 1.304903 | 4 |
| 2 | J | $\alpha$ 1/RMSE | DNN-SO | DNN-SO | EXD3 | 0.983615 | 1.079354 | 1.304903 | 4 |
| S1 | A | Uniform | Any |  | EXD3 | 0.932241 | 1.061235 | 2.153249 | 5 |
| S1 | B | Uniform | All |  | EXD3 | 0.861906 | 0.931562 | 237.3779 | 547 |
| S1 | C | Uniform | LM | BLUP | EXD3 | 0.891821 | 0.895162 | 1.343119 | 4 |
| S1 | C | Uniform | LM | SVR | EXD3 | 0.943201 | 0.986786 | 1.343119 | 4 |
| S1 | C | Uniform | LM | DNN-CO | EXD3 | 0.916316 | 0.921968 | 1.343119 | 4 |
| S1 | C | Uniform | LM | DNN-SO | EXD3 | 0.948092 | 0.969883 | 1.343119 | 4 |
| S1 | C | Uniform | LM | LM | Linear | 0.980282 |  |  |  |
| S1 | C | Uniform | LM | KNN | Linear | 0.980282 |  |  |  |
| S1 | C | Uniform | LM | RNR | Linear | 0.980282 |  |  |  |
| S1 | C | Uniform | LM | RF | Linear | 0.980282 |  |  |  |
| S1 | D | Uniform | BLUP | LM | EXD3 | 0.894109 | 0.89305 | 1.834583 | 5 |
| S1 | D | Uniform | BLUP | BLUP | EXD3 | 0.8943 | 0.989285 | 1.834583 | 5 |
| S1 | D | Uniform | BLUP | KNN | EXD3 | 0.932856 | 0.940406 | 1.834583 | 5 |
| S1 | D | Uniform | BLUP | RNR | EXD3 | 0.944958 | 0.958248 | 1.834583 | 5 |
| S1 | D | Uniform | BLUP | RF | EXD3 | 0.920784 | 0.93151 | 1.834583 | 5 |
| S1 | D | Uniform | BLUP | SVR | EXD3 | 0.886807 | 0.934522 | 1.834583 | 5 |
| S1 | D | Uniform | BLUP | DNN-CO | EXD3 | 0.891809 | 0.913569 | 1.834583 | 5 |
| S1 | D | Uniform | BLUP | DNN-SO | EXD3 | 0.914831 | 0.950725 | 1.834583 | 5 |
| S1 | E | Uniform | KNN | BLUP | EXD3 | 0.933086 | 0.943193 | 1.244419 | 3 |
| S1 | E | Uniform | KNN | SVR | EXD3 | 0.989047 | 1.033769 | 1.244419 | 3 |
| S1 | E | Uniform | KNN | DNN-CO | EXD3 | 0.966101 | 0.973001 | 1.244419 | 3 |
| S1 | E | Uniform | KNN | DNN-SO | EXD3 | 1.004703 | 1.019697 | 1.244419 | 3 |
| S1 | E | Uniform | KNN | LM | Linear | 1.034277 |  |  |  |
| S1 | E | Uniform | KNN | KNN | Linear | 1.034277 |  |  |  |
| S1 | E | Uniform | KNN | RNR | Linear | 1.034277 |  |  |  |
| S1 | E | Uniform | KNN | RF | Linear | 1.034277 |  |  |  |
| S1 | F | Uniform | RNR | BLUP | EXD3 | 0.945056 | 0.958561 | 1.730926 | 4 |
| S1 | F | Uniform | RNR | SVR | EXD3 | 0.997843 | 1.033716 | 1.730926 | 4 |
| S1 | F | Uniform | RNR | DNN-CO | EXD3 | 0.974741 | 0.97829 | 1.730926 | 4 |
| S1 | F | Uniform | RNR | DNN-SO | EXD3 | 1.015509 | 1.035097 | 1.730926 | 4 |
| S1 | F | Uniform | RNR | LM | Linear | 1.045396 |  |  |  |
| S1 | F | Uniform | RNR | KNN | Linear | 1.045396 |  |  |  |
| S1 | F | Uniform | RNR | RNR | Linear | 1.045396 |  |  |  |
| S1 | F | Uniform | RNR | RF | Linear | 1.045396 |  |  |  |
| S1 | G | Uniform | RF | BLUP | EXD3 | 0.921272 | 0.932807 | 1.258276 | 3 |
| S1 | G | Uniform | RF | RF | EXD3 | 1.105697 | 1.109482 | 1.258276 | 3 |
| S1 | G | Uniform | RF | SVR | EXD3 | 0.968986 | 1.013257 | 1.258276 | 3 |
| S1 | G | Uniform | RF | DNN-CO | EXD3 | 0.930602 | 0.936389 | 1.258276 | 3 |
| S1 | G | Uniform | RF | DNN-SO | EXD3 | 0.977434 | 1.002015 | 1.258276 | 3 |
| S1 | G | Uniform | RF | LM | Linear | 1.00957 |  |  |  |
| S1 | G | Uniform | RF | KNN | Linear | 1.00957 |  |  |  |
| S1 | G | Uniform | RF | RNR | Linear | 1.00957 |  |  |  |
| S1 | H | Uniform | SVR | LM | EXD3 | 0.943649 | 0.980078 | 1.493248 | 4 |
| S1 | H | Uniform | SVR | BLUP | EXD3 | 0.887128 | 0.946753 | 1.493248 | 4 |
| S1 | H | Uniform | SVR | KNN | EXD3 | 0.988363 | 1.029836 | 1.493248 | 4 |
| S1 | H | Uniform | SVR | RNR | EXD3 | 0.998254 | 1.038108 | 1.493248 | 4 |
| S1 | H | Uniform | SVR | RF | EXD3 | 0.968717 | 1.006708 | 1.493248 | 4 |
| S1 | H | Uniform | SVR | SVR | EXD3 | 0.956551 | 1.127485 | 1.493248 | 4 |
| S1 | H | Uniform | SVR | DNN-CO | EXD3 | 0.919762 | 0.958588 | 1.493248 | 4 |
| S1 | H | Uniform | SVR | DNN-SO | EXD3 | 0.950156 | 1.019807 | 1.493248 | 4 |
| S1 | I | Uniform | DNN-CO | LM | EXD3 | 0.916347 | 0.920633 | 1.553062 | 4 |
| S1 | I | Uniform | DNN-CO | BLUP | EXD3 | 0.892314 | 0.919199 | 1.553062 | 4 |
| S1 | I | Uniform | DNN-CO | KNN | EXD3 | 0.966017 | 0.971862 | 1.553062 | 4 |
| S1 | I | Uniform | DNN-CO | RNR | EXD3 | 0.9748 | 0.978293 | 1.553062 | 4 |
| S1 | I | Uniform | DNN-CO | RF | EXD3 | 0.930489 | 0.935922 | 1.553062 | 4 |
| S1 | I | Uniform | DNN-CO | SVR | EXD3 | 0.919344 | 0.959656 | 1.553062 | 4 |
| S1 | I | Uniform | DNN-CO | DNN-CO | EXD3 | 0.936628 | 0.950228 | 1.553062 | 4 |
| S1 | I | Uniform | DNN-CO | DNN-SO | EXD3 | 0.941047 | 0.971962 | 1.553062 | 4 |
| S1 | J | Uniform | DNN-SO | LM | EXD3 | 0.948247 | 0.972153 | 1.210296 | 3 |
| S1 | J | Uniform | DNN-SO | BLUP | EXD3 | 0.916149 | 0.96268 | 1.210296 | 3 |
| S1 | J | Uniform | DNN-SO | KNN | EXD3 | 1.004745 | 1.019726 | 1.210296 | 3 |
| S1 | J | Uniform | DNN-SO | RNR | EXD3 | 1.015989 | 1.04183 | 1.210296 | 3 |
| S1 | J | Uniform | DNN-SO | RF | EXD3 | 0.977436 | 1.003236 | 1.210296 | 3 |
| S1 | J | Uniform | DNN-SO | SVR | EXD3 | 0.950525 | 1.036279 | 1.210296 | 3 |
| S1 | J | Uniform | DNN-SO | DNN-CO | EXD3 | 0.941564 | 0.978728 | 1.210296 | 3 |
| S1 | J | Uniform | DNN-SO | DNN-SO | EXD3 | 0.984701 | 1.090478 | 1.210296 | 3 |
| S2 | A | Uniform by Type | Any |  | EXD3 | 0.930523 | 1.090416 | 1.978678 | 5 |
| S2 | B | Uniform by Type | All |  | EXD3 | 0.922008 | 0.931002 | 39.06555 | 90 |
| S2 | C | Uniform by Type | LM | BLUP | EXD3 | 0.890767 | 0.966263 | 0.449368 | 2 |
| S2 | C | Uniform by Type | LM | SVR | EXD3 | 0.945898 | 1.066315 | 0.449368 | 2 |
| S2 | C | Uniform by Type | LM | DNN-CO | EXD3 | 0.916725 | 0.942846 | 0.449368 | 2 |
| S2 | C | Uniform by Type | LM | DNN-SO | EXD3 | 0.948097 | 0.999417 | 0.449368 | 2 |
| S2 | C | Uniform by Type | LM | LM | Linear | 0.980266 |  |  |  |
| S2 | C | Uniform by Type | LM | KNN | Linear | 0.980266 |  |  |  |
| S2 | C | Uniform by Type | LM | RNR | Linear | 0.980266 |  |  |  |
| S2 | C | Uniform by Type | LM | RF | Linear | 0.980266 |  |  |  |
| S2 | D | Uniform by Type | BLUP | LM | EXD3 | 0.890581 | 0.900201 | 1.533774 | 4 |
| S2 | D | Uniform by Type | BLUP | BLUP | EXD3 | 0.90187 | 0.969059 | 1.533774 | 4 |
| S2 | D | Uniform by Type | BLUP | KNN | EXD3 | 0.932455 | 0.960988 | 1.533774 | 4 |
| S2 | D | Uniform by Type | BLUP | RNR | EXD3 | 0.945316 | 0.961781 | 1.533774 | 4 |
| S2 | D | Uniform by Type | BLUP | RF | EXD3 | 0.919811 | 0.950973 | 1.533774 | 4 |
| S2 | D | Uniform by Type | BLUP | SVR | EXD3 | 0.886113 | 0.939232 | 1.533774 | 4 |
| S2 | D | Uniform by Type | BLUP | DNN-CO | EXD3 | 0.891871 | 0.919406 | 1.533774 | 4 |
| S2 | D | Uniform by Type | BLUP | DNN-SO | EXD3 | 0.917102 | 0.950977 | 1.533774 | 4 |
| S2 | E | Uniform by Type | KNN | BLUP | EXD3 | 0.933158 | 0.976572 | 0.969893 | 3 |
| S2 | E | Uniform by Type | KNN | SVR | EXD3 | 0.988927 | 1.059294 | 0.969893 | 3 |
| S2 | E | Uniform by Type | KNN | DNN-CO | EXD3 | 0.966322 | 0.971776 | 0.969893 | 3 |
| S2 | E | Uniform by Type | KNN | DNN-SO | EXD3 | 1.003716 | 1.034374 | 0.969893 | 3 |
| S2 | E | Uniform by Type | KNN | LM | Linear | 1.034273 |  |  |  |
| S2 | E | Uniform by Type | KNN | KNN | Linear | 1.034273 |  |  |  |
| S2 | E | Uniform by Type | KNN | RNR | Linear | 1.034273 |  |  |  |
| S2 | E | Uniform by Type | KNN | RF | Linear | 1.034273 |  |  |  |
| S2 | F | Uniform by Type | RNR | BLUP | EXD3 | 0.945311 | 0.961578 | 1.547408 | 4 |
| S2 | F | Uniform by Type | RNR | SVR | EXD3 | 0.997927 | 1.044351 | 1.547408 | 4 |
| S2 | F | Uniform by Type | RNR | DNN-CO | EXD3 | 0.974545 | 0.981443 | 1.547408 | 4 |
| S2 | F | Uniform by Type | RNR | DNN-SO | EXD3 | 1.015282 | 1.040399 | 1.547408 | 4 |
| S2 | F | Uniform by Type | RNR | LM | Linear | 1.045393 |  |  |  |
| S2 | F | Uniform by Type | RNR | KNN | Linear | 1.045393 |  |  |  |
| S2 | F | Uniform by Type | RNR | RNR | Linear | 1.045393 |  |  |  |
| S2 | F | Uniform by Type | RNR | RF | Linear | 1.045393 |  |  |  |
| S2 | G | Uniform by Type | RF | BLUP | EXD3 | 0.92025 | 0.958728 | 1.199967 | 3 |
| S2 | G | Uniform by Type | RF | RF | EXD3 | 1.105734 | 1.109842 | 1.199967 | 3 |
| S2 | G | Uniform by Type | RF | SVR | EXD3 | 0.969078 | 1.01537 | 1.199967 | 3 |
| S2 | G | Uniform by Type | RF | DNN-CO | EXD3 | 0.930814 | 0.941388 | 1.199967 | 3 |
| S2 | G | Uniform by Type | RF | DNN-SO | EXD3 | 0.977285 | 1.005303 | 1.199967 | 3 |
| S2 | G | Uniform by Type | RF | LM | Linear | 1.009542 |  |  |  |
| S2 | G | Uniform by Type | RF | KNN | Linear | 1.009542 |  |  |  |
| S2 | G | Uniform by Type | RF | RNR | Linear | 1.009542 |  |  |  |
| S2 | H | Uniform by Type | SVR | LM | EXD3 | 0.94533 | 0.980156 | 1.079944 | 3 |
| S2 | H | Uniform by Type | SVR | BLUP | EXD3 | 0.88738 | 0.955746 | 1.079944 | 3 |
| S2 | H | Uniform by Type | SVR | KNN | EXD3 | 0.988695 | 1.051035 | 1.079944 | 3 |
| S2 | H | Uniform by Type | SVR | RNR | EXD3 | 0.998822 | 1.060722 | 1.079944 | 3 |
| S2 | H | Uniform by Type | SVR | RF | EXD3 | 0.969377 | 1.019419 | 1.079944 | 3 |
| S2 | H | Uniform by Type | SVR | SVR | EXD3 | 0.959246 | 1.159809 | 1.079944 | 3 |
| S2 | H | Uniform by Type | SVR | DNN-CO | EXD3 | 0.920202 | 0.983103 | 1.079944 | 3 |
| S2 | H | Uniform by Type | SVR | DNN-SO | EXD3 | 0.949989 | 1.042839 | 1.079944 | 3 |
| S2 | I | Uniform by Type | DNN-CO | LM | EXD3 | 0.916492 | 0.923481 | 1.199138 | 3 |
| S2 | I | Uniform by Type | DNN-CO | BLUP | EXD3 | 0.892282 | 0.924933 | 1.199138 | 3 |
| S2 | I | Uniform by Type | DNN-CO | KNN | EXD3 | 0.966285 | 0.970651 | 1.199138 | 3 |
| S2 | I | Uniform by Type | DNN-CO | RNR | EXD3 | 0.974636 | 0.983381 | 1.199138 | 3 |
| S2 | I | Uniform by Type | DNN-CO | RF | EXD3 | 0.930826 | 0.94111 | 1.199138 | 3 |
| S2 | I | Uniform by Type | DNN-CO | SVR | EXD3 | 0.919923 | 0.976264 | 1.199138 | 3 |
| S2 | I | Uniform by Type | DNN-CO | DNN-CO | EXD3 | 0.936905 | 0.960484 | 1.199138 | 3 |
| S2 | I | Uniform by Type | DNN-CO | DNN-SO | EXD3 | 0.942781 | 0.968458 | 1.199138 | 3 |
| S2 | J | Uniform by Type | DNN-SO | LM | EXD3 | 0.947224 | 0.966114 | 1.230708 | 3 |
| S2 | J | Uniform by Type | DNN-SO | BLUP | EXD3 | 0.917563 | 0.957588 | 1.230708 | 3 |
| S2 | J | Uniform by Type | DNN-SO | KNN | EXD3 | 1.003483 | 1.02762 | 1.230708 | 3 |
| S2 | J | Uniform by Type | DNN-SO | RNR | EXD3 | 1.015724 | 1.044922 | 1.230708 | 3 |
| S2 | J | Uniform by Type | DNN-SO | RF | EXD3 | 0.977245 | 1.004775 | 1.230708 | 3 |
| S2 | J | Uniform by Type | DNN-SO | SVR | EXD3 | 0.949604 | 1.030192 | 1.230708 | 3 |
| S2 | J | Uniform by Type | DNN-SO | DNN-CO | EXD3 | 0.942743 | 0.967876 | 1.230708 | 3 |
| S2 | J | Uniform by Type | DNN-SO | DNN-SO | EXD3 | 0.9851 | 1.077147 | 1.230708 | 3 |
| S3 | A | $\alpha$ 1/s2 | Any | | EXD3 | 0.926334 | 1.059301 | 2.471776 | 6 |
| S3 | B | $\alpha$ 1/s2 | All | | EXD3 | 0.914529 | 0.92564 | 28.40954 | 66 |
| S3 | C | $\alpha$ 1/s2 | LM | BLUP | EXD3 | 0.90595 | 0.91881 | 1.243244 | 3 |
| S3 | C | $\alpha$ 1/s2 | LM | SVR | EXD3 | 0.946769 | 0.968016 | 1.243244 | 3 |
| S3 | C | $\alpha$ 1/s2 | LM | DNN-CO | EXD3 | 0.926638 | 0.933031 | 1.243244 | 3 |
| S3 | C | $\alpha$ 1/s2 | LM | DNN-SO | EXD3 | 0.94804 | 0.959815 | 1.243244 | 3 |
| S3 | C | $\alpha$ 1/s2 | LM | LM | Linear | 0.968727 | | | |
| S3 | C | $\alpha$ 1/s2 | LM | KNN | Linear | 0.968727 | | | |
| S3 | C | $\alpha$ 1/s2 | LM | RNR | Linear | 0.968727 | | | |
| S3 | C | $\alpha$ 1/s2 | LM | RF | Linear | 0.968727 | | | |
| S3 | D | $\alpha$ 1/s2 | BLUP | LM | EXD3 | 0.905863 | 0.916659 | 1.067083 | 3 |
| S3 | D | $\alpha$ 1/s2 | BLUP | BLUP | EXD3 | 0.902887 | 0.929317 | 1.067083 | 3 |
| S3 | D | $\alpha$ 1/s2 | BLUP | KNN | EXD3 | 0.920343 | 0.945742 | 1.067083 | 3 |
| S3 | D | $\alpha$ 1/s2 | BLUP | RNR | EXD3 | 0.927833 | 0.96421 | 1.067083 | 3 |
| S3 | D | $\alpha$ 1/s2 | BLUP | RF | EXD3 | 0.948824 | 0.949702 | 1.067083 | 3 |
| S3 | D | $\alpha$ 1/s2 | BLUP | SVR | EXD3 | 0.887332 | 0.987043 | 1.067083 | 3 |
| S3 | D | $\alpha$ 1/s2 | BLUP | DNN-CO | EXD3 | 0.892461 | 0.917067 | 1.067083 | 3 |
| S3 | D | $\alpha$ 1/s2 | BLUP | DNN-SO | EXD3 | 0.93245 | 0.986102 | 1.067083 | 3 |
| S3 | E | $\alpha$ 1/s2 | KNN | BLUP | EXD3 | 0.919708 | 0.942618 | 1.276945 | 3 |
| S3 | E | $\alpha$ 1/s2 | KNN | SVR | EXD3 | 0.977719 | 1.047781 | 1.276945 | 3 |
| S3 | E | $\alpha$ 1/s2 | KNN | DNN-CO | EXD3 | 0.957247 | 0.985901 | 1.276945 | 3 |
| S3 | E | $\alpha$ 1/s2 | KNN | DNN-SO | EXD3 | 0.991862 | 1.041207 | 1.276945 | 3 |
| S3 | E | $\alpha$ 1/s2 | KNN | LM | Linear | 1.034861 | | | |
| S3 | E | $\alpha$ 1/s2 | KNN | KNN | Linear | 1.034861 | | | |
| S3 | E | $\alpha$ 1/s2 | KNN | RNR | Linear | 1.034861 | | | |
| S3 | E | $\alpha$ 1/s2 | KNN | RF | Linear | 1.034861 | | | |
| S3 | F | $\alpha$ 1/s2 | RNR | BLUP | EXD3 | 0.92795 | 0.971213 | 0.92628 | 3 |
| S3 | F | $\alpha$ 1/s2 | RNR | SVR | EXD3 | 0.987383 | 1.060532 | 0.92628 | 3 |
| S3 | F | $\alpha$ 1/s2 | RNR | DNN-CO | EXD3 | 0.963343 | 1.026669 | 0.92628 | 3 |
| S3 | F | $\alpha$ 1/s2 | RNR | DNN-SO | EXD3 | 0.998166 | 1.052465 | 0.92628 | 3 |
| S3 | F | $\alpha$ 1/s2 | RNR | LM | Linear | 1.042798 | | | |
| S3 | F | $\alpha$ 1/s2 | RNR | KNN | Linear | 1.042798 | | | |
| S3 | F | $\alpha$ 1/s2 | RNR | RNR | Linear | 1.042798 | | | |
| S3 | F | $\alpha$ 1/s2 | RNR | RF | Linear | 1.042798 | | | |
| S3 | G | $\alpha$ 1/s2 | RF | BLUP | EXD3 | 0.9481 | 1.113055 | 0.323343 | 1 |
| S3 | G | $\alpha$ 1/s2 | RF | RF | EXD3 | 1.105846 | 1.144208 | 0.323343 | 1 |
| S3 | G | $\alpha$ 1/s2 | RF | SVR | EXD3 | 0.987191 | 1.169153 | 0.323343 | 1 |
| S3 | G | $\alpha$ 1/s2 | RF | DNN-CO | EXD3 | 0.958601 | 1.142217 | 0.323343 | 1 |
| S3 | G | $\alpha$ 1/s2 | RF | DNN-SO | EXD3 | 0.976312 | 1.114483 | 0.323343 | 1 |
| S3 | G | $\alpha$ 1/s2 | RF | LM | Linear | 1.016935 | | | |
| S3 | G | $\alpha$ 1/s2 | RF | KNN | Linear | 1.016935 | | | |
| S3 | G | $\alpha$ 1/s2 | RF | RNR | Linear | 1.016935 | | | |
| S3 | H | $\alpha 1/s2$ | SVR | LM | EXD3 | 0.946688 | 0.967293 | 1.34548 | 4 |
| S3 | H | $\alpha 1/s2$ | SVR | BLUP | EXD3 | 0.886494 | 0.966389 | 1.34548 | 4 |
| S3 | H | $\alpha 1/s2$ | SVR | KNN | EXD3 | 0.977484 | 1.044667 | 1.34548 | 4 |
| S3 | H | $\alpha 1/s2$ | SVR | RNR | EXD3 | 0.986361 | 1.039301 | 1.34548 | 4 |
| S3 | H | $\alpha 1/s2$ | SVR | RF | EXD3 | 0.985647 | 1.012264 | 1.34548 | 4 |
| S3 | H | $\alpha 1/s2$ | SVR | SVR | EXD3 | 0.956841 | 1.127099 | 1.34548 | 4 |
| S3 | H | $\alpha 1/s2$ | SVR | DNN-CO | EXD3 | 0.920394 | 0.976316 | 1.34548 | 4 |
| S3 | H | $\alpha 1/s2$ | SVR | DNN-SO | EXD3 | 0.956006 | 0.999417 | 1.34548 | 4 |
| S3 | I | $\alpha 1/s2$ | DNN-CO | LM | EXD3 | 0.926971 | 0.932883 | 0.965427 | 3 |
| S3 | I | $\alpha 1/s2$ | DNN-CO | BLUP | EXD3 | 0.892748 | 0.915198 | 0.965427 | 3 |
| S3 | I | $\alpha 1/s2$ | DNN-CO | KNN | EXD3 | 0.957565 | 0.995612 | 0.965427 | 3 |
| S3 | I | $\alpha 1/s2$ | DNN-CO | RNR | EXD3 | 0.963294 | 1.02356 | 0.965427 | 3 |
| S3 | I | $\alpha 1/s2$ | DNN-CO | RF | EXD3 | 0.958191 | 0.981243 | 0.965427 | 3 |
| S3 | I | $\alpha 1/s2$ | DNN-CO | SVR | EXD3 | 0.921268 | 0.997351 | 0.965427 | 3 |
| S3 | I | $\alpha 1/s2$ | DNN-CO | DNN-CO | EXD3 | 0.938532 | 0.974344 | 0.965427 | 3 |
| S3 | I | $\alpha 1/s2$ | DNN-CO | DNN-SO | EXD3 | 0.95283 | 0.996841 | 0.965427 | 3 |
| S3 | J | $\alpha 1/s2$ | DNN-SO | LM | EXD3 | 0.947959 | 0.958787 | 1.396119 | 4 |
| S3 | J | $\alpha 1/s2$ | DNN-SO | BLUP | EXD3 | 0.931846 | 0.974242 | 1.396119 | 4 |
| S3 | J | $\alpha 1/s2$ | DNN-SO | KNN | EXD3 | 0.991614 | 1.037709 | 1.396119 | 4 |
| S3 | J | $\alpha 1/s2$ | DNN-SO | RNR | EXD3 | 0.997414 | 1.034493 | 1.396119 | 4 |
| S3 | J | $\alpha 1/s2$ | DNN-SO | RF | EXD3 | 0.974726 | 0.998323 | 1.396119 | 4 |
| S3 | J | $\alpha 1/s2$ | DNN-SO | SVR | EXD3 | 0.955918 | 0.998104 | 1.396119 | 4 |
| S3 | J | $\alpha 1/s2$ | DNN-SO | DNN-CO | EXD3 | 0.952254 | 0.984409 | 1.396119 | 4 |
| S3 | J | $\alpha 1/s2$ | DNN-SO | DNN-SO | EXD3 | 0.986012 | 1.06255 | 1.396119 | 4 |
| S4 | A | $\alpha 1/s$ | Any | | EXD3 | 0.92909 | 1.067725 | 2.164665 | 5 |
| S4 | B | $\alpha 1/s$ | All | | EXD3 | 0.886797 | 0.927839 | 174.0333 | 401 |
| S4 | C | $\alpha 1/s$ | LM | BLUP | EXD3 | 0.898577 | 0.909271 | 1.448218 | 4 |
| S4 | C | $\alpha 1/s$ | LM | SVR | EXD3 | 0.944187 | 0.975807 | 1.448218 | 4 |
| S4 | C | $\alpha 1/s$ | LM | DNN-CO | EXD3 | 0.921998 | 0.930156 | 1.448218 | 4 |
| S4 | C | $\alpha 1/s$ | LM | DNN-SO | EXD3 | 0.947685 | 0.963359 | 1.448218 | 4 |
| S4 | C | $\alpha 1/s$ | LM | LM | Linear | 0.973201 | | | |
| S4 | C | $\alpha 1/s$ | LM | KNN | Linear | 0.973201 | | | |
| S4 | C | $\alpha 1/s$ | LM | RNR | Linear | 0.973201 | | | |
| S4 | C | $\alpha 1/s$ | LM | RF | Linear | 0.973201 | | | |
| S4 | D | $\alpha 1/s$ | BLUP | LM | EXD3 | 0.898609 | 0.908939 | 1.445051 | 4 |
| S4 | D | $\alpha 1/s$ | BLUP | BLUP | EXD3 | 0.902816 | 0.978671 | 1.445051 | 4 |
| S4 | D | $\alpha 1/s$ | BLUP | KNN | EXD3 | 0.925699 | 0.945227 | 1.445051 | 4 |
| S4 | D | $\alpha 1/s$ | BLUP | RNR | EXD3 | 0.936371 | 0.955168 | 1.445051 | 4 |
| S4 | D | $\alpha 1/s$ | BLUP | RF | EXD3 | 0.934283 | 0.950786 | 1.445051 | 4 |
| S4 | D | $\alpha 1/s$ | BLUP | SVR | EXD3 | 0.887608 | 0.940832 | 1.445051 | 4 |
| S4 | D | $\alpha 1/s$ | BLUP | DNN-CO | EXD3 | 0.891299 | 0.908118 | 1.445051 | 4 |
| S4 | D | $\alpha 1/s$ | BLUP | DNN-SO | EXD3 | 0.921961 | 0.973024 | 1.445051 | 4 |
| S4 | E | $\alpha 1/s$ | KNN | BLUP | EXD3 | 0.925919 | 0.949029 | 1.17696 | 3 |
| S4 | E | $\alpha 1/s$ | KNN | SVR | EXD3 | 0.983748 | 1.051422 | 1.17696 | 3 |
| S4 | E | $\alpha 1/s$ | KNN | DNN-CO | EXD3 | 0.963403 | 0.983445 | 1.17696 | 3 |
| S4 | E | $\alpha 1/s$ | KNN | DNN-SO | EXD3 | 0.996296 | 1.02381 | 1.17696 | 3 |
| S4 | E | $\alpha 1/s$ | KNN | LM | Linear | 1.033501 | | | |
| S4 | E | $\alpha 1/s$ | KNN | KNN | Linear | 1.033501 | | | |
| S4 | E | $\alpha 1/s$ | KNN | RNR | Linear | 1.033501 | | | |
| S4 | E | $\alpha 1/s$ | KNN | RF | Linear | 1.033501 | | | |
| S4 | F | $\alpha 1/s$ | RNR | BLUP | EXD3 | 0.93646 | 0.956491 | 1.335137 | 4 |
| S4 | F | $\alpha 1/s$ | RNR | SVR | EXD3 | 0.992235 | 1.057596 | 1.335137 | 4 |
| S4 | F | $\alpha 1/s$ | RNR | DNN-CO | EXD3 | 0.971636 | 0.986816 | 1.335137 | 4 |
| S4 | F | $\alpha 1/s$ | RNR | DNN-SO | EXD3 | 1.005275 | 1.038688 | 1.335137 | 4 |
| S4 | F | $\alpha 1/s$ | RNR | LM | Linear | 1.042915 | | | |
| S4 | F | $\alpha 1/s$ | RNR | KNN | Linear | 1.042915 | | | |
| S4 | F | $\alpha 1/s$ | RNR | RNR | Linear | 1.042915 | | | |
| S4 | F | $\alpha 1/s$ | RNR | RF | Linear | 1.042915 | | | |
| S4 | G | $\alpha 1/s$ | RF | BLUP | EXD3 | 0.933854 | 0.955749 | 1.457933 | 4 |
| S4 | G | $\alpha 1/s$ | RF | RF | EXD3 | 1.10578 | 1.108244 | 1.457933 | 4 |
| S4 | G | $\alpha 1/s$ | RF | SVR | EXD3 | 0.976776 | 1.00929 | 1.457933 | 4 |
| S4 | G | $\alpha 1/s$ | RF | DNN-CO | EXD3 | 0.946267 | 0.970042 | 1.457933 | 4 |
| S4 | G | $\alpha 1/s$ | RF | DNN-SO | EXD3 | 0.976047 | 0.994761 | 1.457933 | 4 |
| S4 | G | $\alpha 1/s$ | RF | LM | Linear | 1.012001 | | | |
| S4 | G | $\alpha 1/s$ | RF | KNN | Linear | 1.012001 | | | |
| S4 | G | $\alpha 1/s$ | RF | RNR | Linear | 1.012001 | | | |
| S4 | H | $\alpha 1/s$ | SVR | LM | EXD3 | 0.944408 | 0.980634 | 1.234394 | 3 |
| S4 | H | $\alpha 1/s$ | SVR | BLUP | EXD3 | 0.888103 | 0.948407 | 1.234394 | 3 |
| S4 | H | $\alpha 1/s$ | SVR | KNN | EXD3 | 0.983608 | 1.048468 | 1.234394 | 3 |
| S4 | H | $\alpha 1/s$ | SVR | RNR | EXD3 | 0.9925 | 1.062406 | 1.234394 | 3 |
| S4 | H | $\alpha 1/s$ | SVR | RF | EXD3 | 0.977108 | 1.014389 | 1.234394 | 3 |
| S4 | H | $\alpha 1/s$ | SVR | SVR | EXD3 | 0.959889 | 1.137389 | 1.234394 | 3 |
| S4 | H | $\alpha 1/s$ | SVR | DNN-CO | EXD3 | 0.919452 | 0.999854 | 1.234394 | 3 |
| S4 | H | $\alpha 1/s$ | SVR | DNN-SO | EXD3 | 0.953568 | 1.004126 | 1.234394 | 3 |
| S4 | I | $\alpha 1/s$ | DNN-CO | LM | EXD3 | 0.922041 | 0.938973 | 1.042432 | 3 |
| S4 | I | $\alpha 1/s$ | DNN-CO | BLUP | EXD3 | 0.891454 | 0.917139 | 1.042432 | 3 |
| S4 | I | $\alpha 1/s$ | DNN-CO | KNN | EXD3 | 0.963501 | 0.986025 | 1.042432 | 3 |
| S4 | I | $\alpha 1/s$ | DNN-CO | RNR | EXD3 | 0.971836 | 0.990566 | 1.042432 | 3 |
| S4 | I | $\alpha 1/s$ | DNN-CO | RF | EXD3 | 0.946651 | 0.979357 | 1.042432 | 3 |
| S4 | I | $\alpha 1/s$ | DNN-CO | SVR | EXD3 | 0.920031 | 1.014437 | 1.042432 | 3 |
| S4 | I | $\alpha 1/s$ | DNN-CO | DNN-CO | EXD3 | 0.938793 | 0.947056 | 1.042432 | 3 |
| S4 | I | $\alpha 1/s$ | DNN-CO | DNN-SO | EXD3 | 0.947405 | 0.983379 | 1.042432 | 3 |
| S4 | J | $\alpha 1/s$ | DNN-SO | LM | EXD3 | 0.947591 | 0.962529 | 1.565979 | 4 |
| S4 | J | $\alpha 1/s$ | DNN-SO | BLUP | EXD3 | 0.921676 | 0.969991 | 1.565979 | 4 |
| S4 | J | $\alpha 1/s$ | DNN-SO | KNN | EXD3 | 0.995833 | 1.018314 | 1.565979 | 4 |
| S4 | J | $\alpha 1/s$ | DNN-SO | RNR | EXD3 | 1.004959 | 1.034596 | 1.565979 | 4 |
| S4 | J | $\alpha 1/s$ | DNN-SO | RF | EXD3 | 0.97557 | 0.997908 | 1.565979 | 4 |
| S4 | J | $\alpha 1/s$ | DNN-SO | SVR | EXD3 | 0.95285 | 0.995468 | 1.565979 | 4 |
| S4 | J | $\alpha 1/s$ | DNN-SO | DNN-CO | EXD3 | 0.946845 | 0.973133 | 1.565979 | 4 |
| S4 | J | $\alpha 1/s$ | DNN-SO | DNN-SO | EXD3 | 0.984062 | 1.066558 | 1.565979 | 4 |

**Supplementary Table 3.** Genetic Algorithm Optimized Ensembling Strategy The best ten combinations of model replicates (selected at random with replacement) and ensembling strategy as optimized though a genetic algorithm.

| Ensemble | LM | BLUP | KNN | RF | RNR | SVR | DNN-SO | DNN-CO | RMSE |
| --- | --- | --- | --- | --- | --- | --- | --- | --- | --- |
| $\alpha$ 1/RMSE | 2 | 8 | 0 | 4 | 0 | 2 | 0 | 6 | 0.8723 |
| $\alpha$ 1/RMSE | 6 | 8 | 0 | 3 | 0 | 3 | 0 | 1 | 0.872691 |
| $\alpha$ 1/RMSE | 6 | 8 | 0 | 2 | 0 | 3 | 0 | 1 | 0.873484 |
| $\alpha$ 1/RMSE | 2 | 8 | 0 | 3 | 0 | 2 | 0 | 6 | 0.874447 |
| $\alpha$ 1/RMSE | 3 | 8 | 0 | 3 | 0 | 2 | 0 | 6 | 0.874618 |
| $\alpha$ 1/RMSE | 2 | 8 | 0 | 3 | 0 | 3 | 0 | 4 | 0.874649 |
| $\alpha$ 1/RMSE | 2 | 8 | 0 | 4 | 0 | 6 | 0 | 4 | 0.874961 |
| $\alpha$ 1/RMSE | 6 | 8 | 0 | 3 | 0 | 2 | 0 | 6 | 0.875245 |
| $\alpha$ 1/RMSE | 5 | 8 | 0 | 4 | 0 | 2 | 0 | 6 | 0.875298 |
| $\alpha$ 1/RMSE | 2 | 8 | 0 | 1 | 0 | 6 | 0 | 4 | 0.875309 |

